# Personalized Risk Prediction for Cancer Survivors: A Bayesian Semi-parametric Recurrent Event Model with Competing Outcomes

**DOI:** 10.1101/2023.02.28.530537

**Authors:** Nam H Nguyen, Seung Jun Shin, Elissa B Dodd-Eaton, Jing Ning, Wenyi Wang

## Abstract

Multiple primary cancers are increasingly more frequent due to improved survival of cancer patients. Characteristics of the first primary cancer largely impact the risk of developing subsequent primary cancers. Hence, model-based risk characterization of cancer survivors that captures patient-specific variables is needed for healthcare policy making. We propose a Bayesian semi-parametric framework, where the occurrence processes of the competing cancer types follow independent non-homogeneous Poisson processes and adjust for covariates including the type and age at diagnosis of the first primary. Applying this framework to a historically collected cohort with families presenting a highly enriched history of multiple primary tumors and diverse cancer types, we have derived a suite of age-to-onset penetrance curves for cancer survivors. This includes penetrance estimates for second primary lung cancer, potentially impactful to ongoing cancer screening decisions. Using Receiver Operating Characteristic (ROC) curves, we have validated the good predictive performance of our models in predicting second primary lung cancer, sarcoma, breast cancer, and all other cancers combined, with areas under the curves (AUCs) at 0.89, 0.91, 0.76 and 0.68, respectively. In conclusion, our framework provides covariate-adjusted quantitative risk assessment for cancer survivors, hence moving a step closer to personalized health management for this unique population.

## 1 Introduction

The most recent report from the National Cancer Institute’s Surveillance, Epidemiology, and End Results (SEER) program estimated approximately 20% of all incident cancer cases in the United States are second or later primary cancers, i.e., additional cancers occurring in cancer survivors (Forjaz et al., 2022). By 2026, the total number of cancer survivors living in the US will exceed 20 million (Hulvat, 2020). Currently, the health care management of cancer survivors is largely indifferent from the management of either at-risk healthy individuals or new cancer patients, thus not yet adequately configured to deliver the desirable high-quality care in this growing population (Mayer et al., 2017). Recently, large population-based epidemiology studies have identified a few important features for cancer survivors: patients with bladder cancer present the highest risk of developing second cancer (Donin et al., 2016); and patients with breast cancer have a higher risk of developing a second primary cancer, lung cancer in particular, than the general population (Bao et al., 2021). Accurate characterization of the onset of second primary cancers is essential for cancer prevention strategy development, such as the United States Preventive Services Task Force (USPSTF) guidelines. One such example is the lung-cancer screening, which has been recommended to adults who are deemed at high-risk, because of the significant survival improvement with diagnosis at an early stage. However, the current recommendation does not include previous malignancy as a high-risk feature (Nobel et al., 2022), due to a lack of accurate personalized risk characterization in such a population.

The NCI SEER program provides a range of databases that allow researchers to directly query cancer-specific incidence rates, defined as the number of new cases per 100,000 people per year, for the first primary cancer (www.seer.cancer.gov/seerstat version 8.4.0.1). These incidence rates can be segmented by 5-year age groups between age 0 and 85, as well as by basic demographic features such as gender. To address the increasing number of cancer survivors, the program recently updated its SEER*Stat software to allow for queries from patients that developed a specific cancer type as the second primary. Although the incidence rates for the second primary cancer can be approximated from the SEER data, such approximation is subject to biases, and hence does not accurately reflect age-at-onset penetrances. Furthermore, these rates are not fully personalized, i.e., they do not account for important person-specific features beyond age group and gender. For example, in the context of inherited cancer syndromes, where the personal risk of developing cancer is significantly increased by the carrier status of a genetic mutation (Parmigiani et al., 1998; Chen et al., 2006), the necessary covariate of mutation status is not captured by SEER.

Although covariate-adjusted risk characterization is urgently needed, unbiased and comprehensive data collection for cancer survivors, as needed for mathematical modeling, remains a major bottleneck. So far, a pan-cancer risk characterization study is not available except for the SEER study mentioned above. On the other hand, there exists a well-characterized and well-ascertained inherited and rare cancer syndrome, called Li-Fraumeni syndrome (LFS) (Li and Fraumeni, 1969), which is mainly caused by germline mutations in the tumor suppressor gene *TP53* (Malkin et al., 1990). LFS presents a wide spectrum of cancer types, as well as a much higher incidence rate of multiple primary cancers than the general population (50% as compared to 2-17%) (Mai et al., 2016; Vogt et al., 2017). We therefore resort to datasets that were collected from patients affected by LFS (see **Table 1** for a dataset example) for real-data motivation, model training and the model-based risk prediction application. People affected by LFS are more likely to develop a spectrum of cancer types, including breast cancer, soft-tissue sarcoma, osteosarcoma, and others, and repeatedly over their lifetime (Li and Fraumeni, 1969). Because of the inheritable nature of *TP53* mutations, the family members of patients diagnosed with LFS and the patients themselves live with a constant high level of anxiety, thus actively participating in risk counseling and cancer screening. This further highlights the need for cancer-specific risk prediction among cancer survivors while addressing genetic mutations as covariates.

**Table 1:** Categorization of family members in the MDACC cohort by gender, number of primary cancers and *TP53* mutation status. FPC: first primary cancer; SPC: second primary cancer. Families are grouped into two categories, positive: at least one family member with a *TP53* mutation, and negative: no mutation carriers.

|  | Positive families |  |  |  | Negative families |  |  |  |
| --- | --- | --- | --- | --- | --- | --- | --- | --- |
|  | Wildtype | Mutation | Unknown | Total | Wildtype | Mutation | Unknown | Total |
| **Male** |  |  |  |  |  |  |  |  |
| Healthy | 112 | 41 | 2,227 | **2,380** | 7 | 0 | 2,221 | **2,228** |
| FPC | 11 | 50 | 339 | **400** | 47 | 0 | 408 | **455** |
| SPC | 1 | 39 | 17 | **57** | 32 | 0 | 43 | **75** |
| Subtotal | 124 | 130 | 2,583 | **2,837** | 86 | 0 | 2,672 | **2,758** |
| **Female** |  |  |  |  |  |  |  |  |
| Healthy | 133 | 32 | 1,997 | **2,162** | 11 | 0 | 2,119 | **2,130** |
| FPC | 9 | 76 | 375 | **460** | 104 | 0 | 423 | **527** |
| SPC | 1 | 97 | 41 | **139** | 113 | 0 | 60 | **173** |
| Subtotal | 143 | 205 | 2,413 | **2,761** | 228 | 0 | 2,602 | **2,830** |
| **Total** | 267 | 335 | 4,996 | **5,598** | 314 | 0 | 5,274 | **5,588** |

Other than risk modeling for individual data, we also take the opportunity to develop additional methods that address the family inheritance structure presented in the LFS datasets. There are several models in the literature that attempt personalized risk prediction given family data. Chen et al. (2009) proposed a frailty model with two-phase case-control data. Choi (2012) introduced a shared frailty model for analyzing correlated time-to-event data arising from family-based studies. Both models, however, do not appropriately account for the pedigree structure of families that is needed for genetic inheritance models. Choi et al. (2016) took one step further by proposing a progressive three-state Markov model, in which a single-step EM algorithm is used to calculate genotype probabilities for individuals with no genetic testing results based on genotype data from other family members. Shin et al. (2019) and Shin et al. (2020) developed Bayesian semiparametric models, which introduced the peeling algorithm (Elston and Stewart, 1971) to take into account familial genetic structure, and the ascertainment-corrected joint (ACJ) approach (Iversen and Chen, 2005) to correct for ascertainment bias. However, none of these approaches have been able to model multiple cancer types beyond the first primary cancer.

In this paper, we introduce a novel Bayesian semiparametric framework that jointly models both multiple primary cancers and multiple cancer types. We utilize non-homogeneous Poisson processes to model the occurrence processes of primary cancers, each of which is characterized by an intensity function that is cancer-type-specific. This modeling approach allows predictions of the second primary to be dependent on the type and timing of the first primary. Under this novel framework, we derive an explicit expression for the individual likelihood contribution, and further a family-wise likelihood through integration across family members. We train and cross-validate our model on a patient cohort, collected from MD Anderson Cancer Center (MDACC) using clinical LFS criteria (Li et al., 1988; Chompret et al., 2001; Bougeard et al., 2015) from year 2000 to 2015 (**Table 1**).

The paper is organized as follows. In Section 2, we present the modeling and derivation of individual and family-wise likelihoods, techniques for addressing necessary ascertainment bias correction and incorporating frailty into family data, as well as simulation studies to evaluate the parameter estimation performances. In Section 3, we demonstrate how the individual likelihood can be used to construct effective risk models, with a special emphasis on lung cancer. In Section 4, we apply our family-wise model to the LFS dataset to obtain novel penetrance estimates for *TP53* mutation carriers and non-carriers. Both sections include a cross-validation study. Section 5 concludes with future research directions.

## 2 Method

### 2.1 Age-at-onset penetrance of multiple cancer types in multiple primaries

We assume multiple cancer occurrences as recurrent events (Cook and Lawless, 2007). To describe the model, we begin by introducing some notations. Let *N*(*t*) be a non-homogeneous Poisson process that counts the number of primary cancers by age *t*, and *Z* be the indicator of cancer type (i.e. *Z* ∈ {1,…, *K*}) with *K* being the number of cancer types. In addition, *H*(*t*) denotes the historical information of *N*(*t*) up to time *t*, and ***X*** is a vector of patient-specific covariates, respectively. The kth cancer-specific intensity function, *λ_k_*(*t*|**X**, *H*(*t*)), *k* =1, 2, …, *K*, is defined as

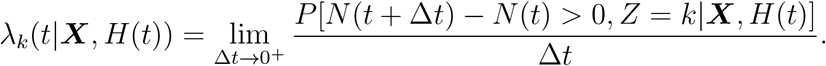

We note that the overall intensity of *N*(*t*) is 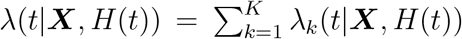. The notion of age-at-onset penetrance is defined as the probability that a patient develops the disease of interest by age t given his or her characteristics (Langbehn et al., 2004). Since then, the definition has been extended to accommodate more complex settings. For example, the *k*th cancer-specific age-at-onset penetrance is defined as the conditional probability of having the *k*th type of cancer by age *t* prior to developing other cancer types. Compared to our previous model that estimates age-at-onset cancer-specific penetrance among the first primary cancers (Shin et al., 2020), here we focus on addressing the need of characterizing the *k*th cancer type among multiple primary cancers during the patient’s lifetime. The desired solution will add another layer of complexity to further extend the definition of penetrance. To this end, let *L* be a non-negative integer that denotes the number of primary cancer occurrences. Then, we define the cancer-specific age-at-onset penetrance for multiple primary cancer as the probability of having the kth cancer type at the *l*th occurrence by age *t* given the (*l* – 1)th occurrence at time *u* in the past, conditional on covariates representing patient characteristics and cancer history up to a time point *u*. Let *T_l_* be the time and *Z_l_* be the cancer type of the *l*th cancer diagnosis. The penetrance of interest, denoted by *q_kl_*(*t*|*u*, ***X***(*u*)), *k* = 1,…, *K*; *l* = 1, …, *L*, is then given by

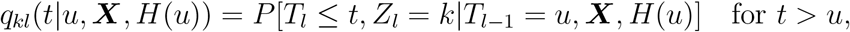

where *T*_0_ = 0 for convenience. For *t* > *u*, we then have

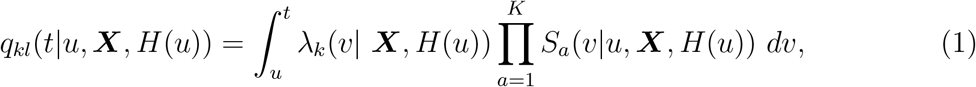

where 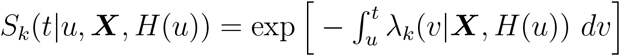 for *k* =1, …,*K*.

For a patient with *l* – 1 primary cancers, we are also interested in the probability of having the next cancer of any type by age *t* given the (*l* – 1)th occurrence at time *u*. This penetrance can be obtained by summing over all cancer types, i.e., 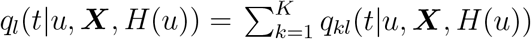.

### 2.2 Intensity function modeling

One of the challenges in modeling recurrent events with competing risks is accounting for the correlations between the events. To address this, we introduce a binary variable *D_k_*(*t*) that indicates whether the patient had the *k*th cancer type as the first primary by time *t*. That is, *D_k_*(*t*) = 1 if *T*_1_ < *t* and *Z*_1_ = *k*, and *D_k_*(*t*) = 0 otherwise. The variable *D_k_*(*t*) allows the risk of later primary to depend on the type of the first one. The relationship between the first and second primary has been established in a number of epidemiological studies (Travis et al., 2005; Mariotto et al., 2007; Donin et al., 2016). We define *D_k,l_*(*t*), *k* = 1,…, *K*, as indicators of the *l*th primary cancer. That is, *D_k,l_*(*t*) = 1 if *T_l_* < *t* and *Z_l_* = *k*. By incorporating the set of covariates {*D_k,l_* (*t*): *k* = 1…, *K* and *l* = 1,…, *L* – 1} into the model specification, we allow the risk of the *l*th primary cancer to depend on the types of the previous *l* – 1 occurrences. In our application, we consider only dependence on the type of the first primary cancer due to limited data availability. For each individual with an inherited cancer syndrome such as LFS, we let *G* be the mutation status (0 for wildtype, 1 for mutated) for the gene of interest, and *S* be the sex (0 for female, 1 for male). Therefore, the covariate vector is ***X***(*t*) = {*G, S, D*_1_(*t*),…, *D_K_*(*t*)}^*T*^.

We model the cancer-specific intensities using a proportional intensity model as follows:

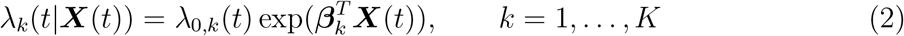

where ***β***_*k*_ denotes the vector of coefficients, and *λ*_0,*k*_(*t*) is the baseline intensity function. Let 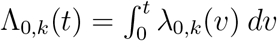 be the cumulative baseline intensity function. We model Λ_0,*k*_ (*t*) using Bernstein polynomials, which are often used to approximate functions with constraints such as monotonicity (Lorentz, 1953; Shin et al., 2020). After rescaling *t* to [0,1], we can approximate Λ_0,*k*_(*t*) using the Bernstein polynomials as follows.

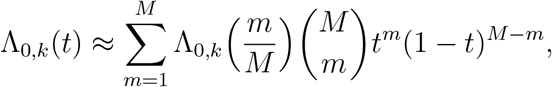

where *M* denotes the order of the Bernstein polynomials. By reparameterizing 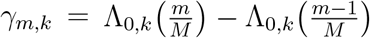 with Λ_0,*k*_(0) = 0, it can be shown that

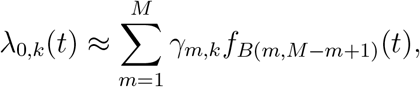

where *f*_*B*(*m,M*–*m*+1)_ denotes the beta density with parameters *m* and *M* – *m* + 1 (Curtis and Ghosh, 2011). In practice, death from other causes is another competing risk that should be taken into account. One can model the hazard function for death in an identical way to the other cancer-specific intensities from (2).

### 2.3 Individual likelihood with known G

We first derive the full likelihood of an individual’s observed cancer history, including the development of first, second, or more primary cancers over time, until censorship, when the genotype *G* is known. For individual *j*, where *j* = 1,2, …, *n*, let *L_j_* be the number of observed cancer occurrences. Although we focus on *L_j_* ≤ 2 in the applications, we describe our model in a more general setting. We observe a dataset 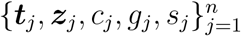, where ***t***_*j*_ and ***z***_*j*_ are *L_j_*-dimensional vectors that indicate the times and cancer types at diagnoses; *c_j_* is the censoring time; and *g_j_* and *s_j_* denote *TP53* mutation status and sex, respectively. Now, we introduce *d_j,k_*(*t*) = 1 if *j* < *t* and *z*_*j*,1_ = *k*, and *d_j,k_*(*t*) = 0 otherwise. Finally, the vector of covariates is given by 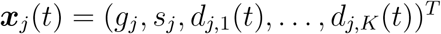.

Let ***β*** = {***β***_*k*_: *k* = 1,…,*K*} and ***γ*** = {*γ_m,k_*: *m* = 1 …,*M*; *k* = 1 …,*K*}. For an individual with cancer history ***h***_*j*_ = {***t***_*j*_, ***z***_*j*_, *c_j_*}, the likelihood can be expressed as a product of the observed cancer occurrences and the censoring event

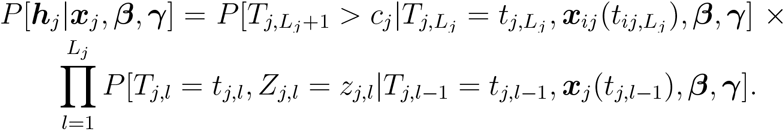

By noting that *d_k_* (*t*), *k* = 1,…, *K*, are periodically fixed, we obtain the likelihood contribution from the lth cancer occurrence as follows.

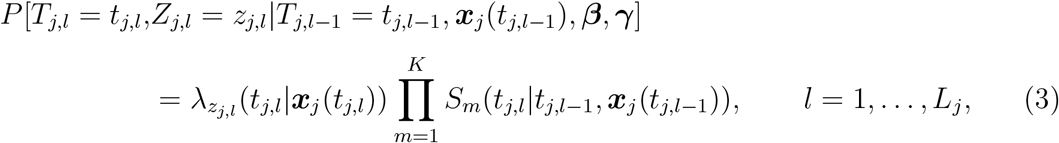

with *t*_*j*,0_ = 0. The likelihood contribution from the censored event is given by

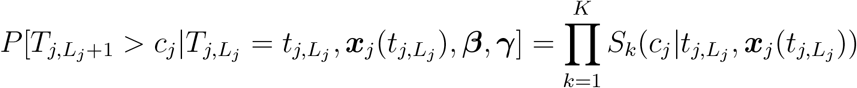

Finally, the likelihood for the *j*th individual is

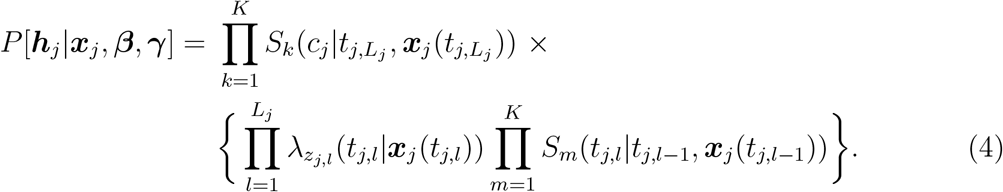

Since we assume the individuals are independent, the overall likelihood of the dataset is simply given by the product of all the individual likelihoods. The complete derivations of the equations above are relegated to **Supplementary Material Section A**.

### 2.4 Family-wise likelihood with unknown *G*

For inherited cancer syndromes in the general population, the total number of carriers is very low, e.g., ~1 out of 1,300 for *TP53* mutations (Gao et al., 2020). This rare condition means even one of the most extensively collected datasets would still have just a few hundred mutation carriers (*n* = 335, see **Table 1**) to be used for estimating a series of penetrance curves, in order to capture dynamic cancer outcomes within the first and the second primary cancers (referred to as SPC and FPC from now on). On the other hand, data collection for inherited cancer syndromes, such as LFS, usually includes cancer history information from family members. Accordingly, we have data from an additional tens of thousands of family members (e.g., *n* = 10, 270, see **Table 1**), who have all the required information except for mutation status. Many of these family members can carry the mutation due to their genetic relationship with the carriers. Therefore, we introduce a model that includes all family members, including those with unknown *G*.

#### 2.4.1 Frailty modeling for the family units

Let *i* = {1,…, *I*} denote the families. In order to account for often-observed non-genetic correlations between family members, after conditioning on ***X***(*t*), as a random effect, we introduce a frailty term (Hougaard, 1995), denoted by *ξ_i,k_*, into the modeling of the cancer-specific intensities (2). Importantly, this family-wise random effect may vary across families and across different cancer types. Hence we have

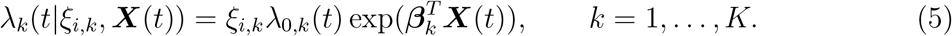

For each cancer type *k*, we assume the frailty term to follow a gamma distribution with the same shape and scale parameters, i.e., 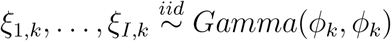. With the presence of the frailties, the age-at-onset penetrance defined in eq.(1) is further modified by marginalizing over the frailties ***ξ*** = {*ξ*_1_,…, *ξ_K_*}. For *t* > *u*, we then have

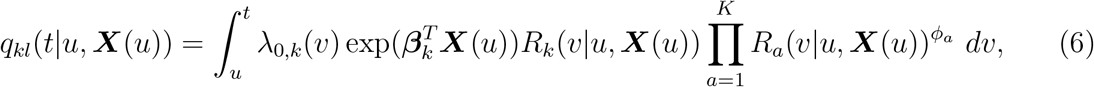

where 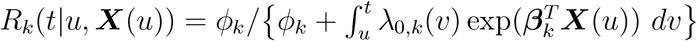.

#### 2.4.2 Peeling algorithm to model the Mendelian inheritance

Let ***θ*** = {***β, γ***, *ϕ*} be the full set of model parameters. The likelihood contribution of the *j*th individual in the ith family, which we call family-wise likelihood, can be derived similarly to (4), when the covariate vector ***x***_*ij*_ is fully observed for all family member *j* in family *i*:

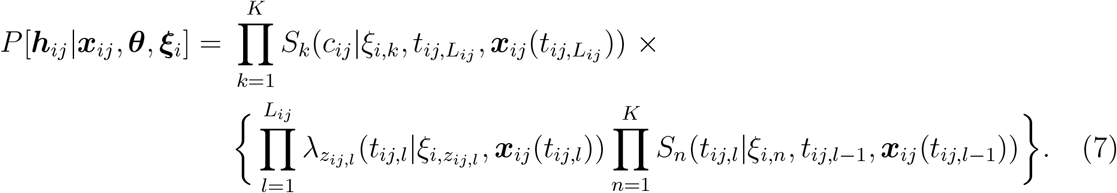

Let ***x***_*i*_ = {***x***_*ij*_: *j* = 1,…,*n_i_* and ***h***_*j*_ = {***h***_*ij*_: *j* = 1,…,*n_i_* be the aggregated covariate vector and cancer history across all family members of the ith family, with ***h***_*ij*_ = {***t***_*ij*_, ***z***_*ij*_, *c_ij_*}. Direct evaluation of (7), however, is not possible in our application since the *TP53* mutation status is unknown for most family members.

To tackle this, we employ the peeling algorithm (Elston and Stewart, 1971) as described in the following. Let ***g***_*i*_ = {*g*_*ij*_: *j* = 1,…,*n_i_*} be the set of genotype information. We then partition the covariate vector into genotype and other non-genetic covariates, hence 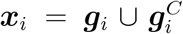. Let ***g***_*i,obs*_ = {*g_ij_*: *g_ij_* is known} be the observed part and ***g**_i,mis_* the missing part of the genotype data for the ith family. We then have ***g**_i_* = ***g***_*i,mis*_ ∪ ***g***_*i,obs*_. We sum over all possible genotype configurations for the family members who have missing genotype information to calculate the family-wise likelihood as 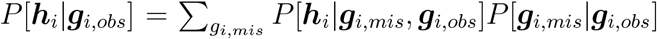. Although this summation procedure seems computationally intractable as the family size increases, the peeling algorithm can solve it very efficiently(Shin et al., 2019, 2020). The details of the implementation of the peeling algorithm are provided in **Supplementary Material Section B**.

#### 2.4.3 Ascertainment bias correction for families with LFS

The ascertainment bias is inevitable and its correction is essential in studying rare diseases like LFS since the data can only be collected from high-risk populations. Introducing an ascertainment indicator for the ith family 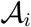 that takes 1 if the ith family is ascertained and 0 otherwise, the ascertainment-corrected joint (ACJ) likelihood (Iversen and Chen, 2005) is

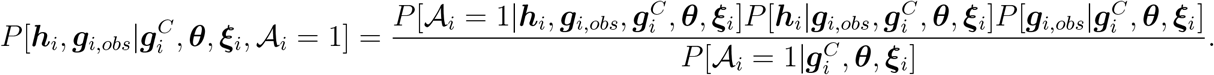

In practice, the ascertainment decision is often made based on the phenotype of the proband in a deterministic way. If so, the first term in the numerator, 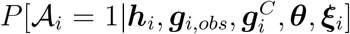 is reduced to 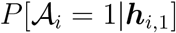 which is independent of the model parameters. In addition, it is not practical for any model to predict genotype data from non-genetic covariates, hence we can reasonably assume that 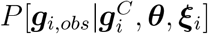 is simplified to *P* [***g***_*i,obs*_]. We then have the following ACJ likelihood up to a constant of proportionality

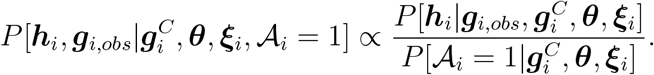

Note that the numerator is the likelihood contribution of the ith family without considering the ascertainment bias, which can be obtained by the peeling algorithm as described in Section 2.4.2. For most LFS studies, the probands can be ascertained due to the diagnosis of a variety of cancer types, in particular those that belong to the LFS spectrum. The ascertainment probability for LFS studies, where the ascertainment decision is based on the first primary cancer diagnosis of the proband, is given by

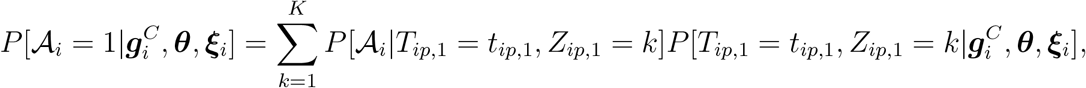

where the subscript *p* denotes the proband. We are interested in the ACJ likelihood up to proportionality, hence only the ratio between the terms 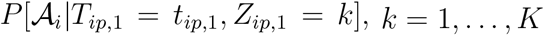, matters. Denoting these terms by *a*_1_,…, *a_K_*, where *a_K_* = 1, it follows that

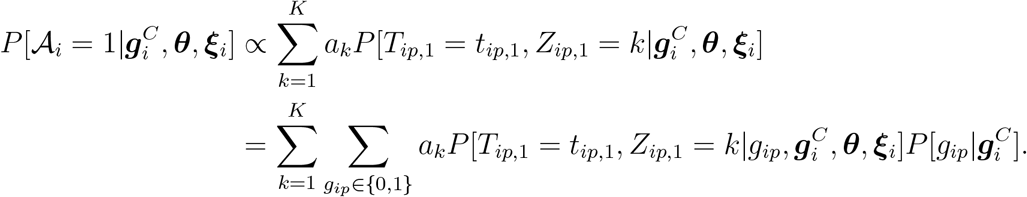

In our application, the *TP53* mutation prevalence is known to be independent of the non-genetic covariates which yields 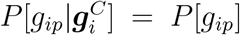. This probability can be calculated readily from the mutated allele frequency *κ_A_*: *P*[*G_ip_* = 0] = (1 – *κ_A_*)^2^, and *P*[*G_ip_* = 1] = 1 – (1 – *κ_A_*)^2^. In the Western population, we have = 0.0006 (Lalloo et al., 2003). In Section 4, we will show that our model is not sensitive to the choice of *a_k_, k* = 1,…, *K*.

We denote the overall likelihood of the dataset by 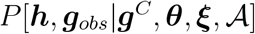, where the family subscript i is dropped to represent aggregation across all the *I* families (e.g., ***h*** = {***h***_*i*_: *i* = 1, …, *I*}). Once the ACJ likelihood has been obtained for each family, the overall likelihood can easily be evaluated as a product as follows, since the families are independent

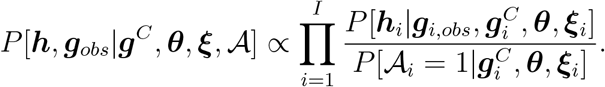

### 2.5 Markov chain Monte Carlo (MCMC) estimation

We assume a normal prior distribution for ***β***_*k*_: ***β***_*k*_ ∼ *N* (**0,σ**^2^***I***), *k* = 1,…, *K*, where **0** and ***I*** denote the zero vector and the identity matrix respectively, and *σ* is set to be large enough (e.g., 100) to reflect little prior knowledge about ***β***_*k*_. For the baseline intensity function, we set *γ_m,k_* ~ *Gamma*(0.01, 0.01), *m* = 1,…,*M* and *k* = 1,…,*K*. This distribution is non-negative as required of *γ_m,k_*, and the corresponding variance is 100 to reflect little prior knowledge. We assume the same gamma prior for *φ_k_, k* = 1,…, *K*. It has been shown previously that the penetrance estimates for real data with *TP53* mutations are not particularly sensitive to the choice of parameter for the gamma prior through a sensitivity analysis (Shin et al., 2020). We denote the priors of ***β, γ***, and ***φ*** by *P*[***β***], *P*[***γ***], and *P*[***φ***], respectively. Then, the joint posterior distribution of ***θ*** and ***ξ*** is given by

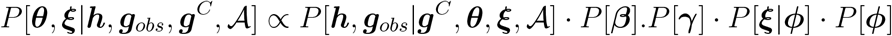

We use a random walk Metropolis-Hastings-within-Gibbs algorithm to generate posterior samples, of which the first part is discarded as burn-in. The MCMC is coded in `R` with the peeling algorithm implemented in `C++` and linked via the `Rcpp` library.

### 2.6 Simulation study

To test the individual likelihood model, we simulated 20 datasets, each consisting of 10,000 individuals with known genotypes. We generated the cancer outcome data for these individuals based on a given set of model parameters with *K* = 2. The MCMC samples are converged well and the corresponding 95% credible intervals cover corresponding true parameters In addition, the estimates show promising performance with low bias and mean squared errors. We refer to **Supplementary Materials Section C** for complete details about the simulation for the individual likelihood model.

For the family-wise likelihood model in Section 2.4, we simulated 20 datasets, each consisting of 300 families with 30 family members spanning three generations. We generated the mutation and the cancer outcome data for all family members, as well as an ascertainment scheme to collect families that meet certain criteria until the total number reaches 300. We include the frailty term and apply the ACJ likelihood for the bias correction. Similar to the individual likelihood case, MCMC estimates converge very well with promising performance. The ***β^G^*** parameters showed the highest absolute biases, 0.35 for 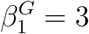 and 0.36 for 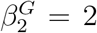. Considering the high complexity of the family-based likelihood model, we are satisfied with its overall performance. **Supplementary Materials Section D** provides full details about the simulation for the family-wise likelihood model.

## 3 Application of the individual likelihood

We apply our models to the MDACC patient cohort as introduced in Section 1. We refer to **Table E.1** (**Supplementary Materials Section E**) for more details about about the first and second primary cancers in the dataset. Since the positive families have characteristics that are distinct from the negative ones (e.g., more mutation carriers, more family members with SPC), we put these two types of families into two separate strata. For the crossvalidation study, we split the full dataset, randomly with each strata, into two subsets of equal size to have training and validation datasets comparable in cancer-type distributions. See **Table E.2** (**Supplementary Materials Section E**).

To apply the individual likelihood model to individuals with known G in the training set, we focused on all family members who were tested for *TP53*, plus those with a predicted mutation probability greater than 0.9 (inferred mutation carriers) or less than 0.01 (inferred non-carriers). These probabilities were calculated using the peeling algorithm as described in Section 2.4.2. To utilize the individual likelihood-based penetrance estimates in the validation set, we expanded our validation subjects to include family members with a predicted mutation probability greater than 0.9 or less than 0.01. Related summaries of the training and validation sets for the individual likelihood model are provided in **Tables E.3 and E.4** (**Supplementary Materials Section E**).

We also compare our penetrance estimates with those that are imputed from the most recent SEER Research Plus database (2022). The database provides access to incidence data across eight registries in the US from 1975 to 2019. See **Table E.5** (**Supplementary Materials Section E**) for the summary of the SEER data.

### 3.1 Model specifications

We choose three individual cancer-type models to demonstrate utility, with each focused on a different cancer type group among the second primary cancers: i) breast cancer, ii) sarcoma, and iii) lung cancer. Here we group all the other cancer types in contrast with the chosen one to maximize sample size and reduce the number of parameters to be estimated. Our model design is based on the frequency of cancer types among SPC (**Tables E.1 – E.4, Supplementary Materials Section E**) as well as the potential relevance to public health (e.g., lung cancer screening). In each model, we categorize the outcomes as breast cancer (or sarcoma, lung cancer), other cancers, and death. For clarity of notations, we let *k* ∈ {*br/sa/lu, ot, d*}, instead of integers, where *br, sa, lu, ot*, and *d* refer to breast cancer, sarcoma, lung cancer, other cancers, and death respectively. For the latter two groups, the intensity/hazard functions take the form given by Equation (4), where **X**(*t*) = {*G,S, D*(*t*)}^*T*^, and *D*(*t*) is an indicator of previous cancer occurrences (i.e., *D*(*t*) = 1 if the person has had at least one cancer in the past, and *D*(*t*) = 0 otherwise). Here we use a common indicator to alleviate limited data availability, but cancer-specific indicators, as used in the simulation study, is still applicable to obtain insights on interactions between cancer types given enough data. Given this covariate vector ***X*** (*t*), we write 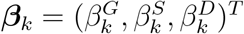 for the regression coefficients in the intensity of cancer type *k*. Our dataset has no male patients diagnosed with breast cancer. Hence we restrict the intensity of breast cancer to 0 for male as 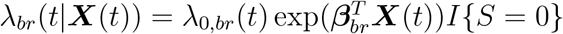. The sarcoma model categorizes the outcomes into sarcoma, other cancers, and death (i.e., *k* ∈ {*sa,ot,d*}). Lastly, the lung cancer model considers lung cancer, other cancers, and death (i.e., *k* ∈ {*lu, ot, d*}). For both models, we do not impose any restrictions such as the indicator function above on their intensity functions.

### 3.2 Model estimation

Using the random-walk Metropolis-Hastings-within-Gibbs algorithm, we obtain 10,000 posterior samples, of which the first 5,000 are discarded as burn-in. Each MCMC iteration takes about two minutes on a high-performance computing cluster at MDACC (407 nodes, each of which has 28 CPU cores with Intel Skylake 2.6Ghz). Posterior samples for all model parameters converged well within the first 10,000 iterations (**Figures F.1–F.3, Supplementary Material Section F**).

**Table F.4** (**Supplementary Material Section F**) shows the estimated parameters. We observe that 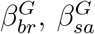, and 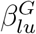 are significantly positive across all models as expected, since breast cancer, sarcoma, and lung cancer all belong to the LFS spectrum. The parameter *βde^D^* is also significantly positive in all three models as patients with one cancer in the past are at higher risk of mortality. In the lung cancer model, it is interesting to see that 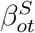 is significantly negative, suggesting that females are more likely to develop cancers outside of the lung. In light of our competing risk framework, it may be due to the prevalence of breast cancer, which is exclusive to females in our training data, within the group of other cancer types. This hypothesis is further confirmed by observing that 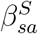 in the sarcoma model is not significant, indicating that sarcoma, the second most popular cancer type in LFS, does not present gender disparity.

### 3.3 Age-at-onset penetrance

For each of the three models in Section 3.1, our primary goal is to estimate the cancer-specific age-at-onset penetrances. Using the notations introduced in Section 2, for a patient with *TP53* mutation status g and sex s, the set of cancer-specific penetrances to the first primary is denoted by *q*_*k*1_(*t*|0, ***X***), *k* = 1, 2, 3, where ***X*** = {*g,s*, 0}^*T*^. For a patient with a first primary of any type at age 10, the set of cancer-specific penetrances to the second primary is given by *q*_*k*2_(*t*|10, ***X***), *k* = 1,2, 3, where ***X*** = {*g, s*, 1}^*T*^. While we can construct penetrance curves for other cancers and death, the primary purpose of the Breast Cancer model is to estimate penetrance for breast cancer (e.g., *q*_*br*,1_,(*t*|0, ***X***) and *q*_*br*,2_(*t*|10, ***X***)), which is of clinical relevance. Similarly, we estimate the age-at-onset penetrances to the first and second primary for sarcoma, and lung cancer, respectively.

The estimated age-at-onset penetrance to the first primary is presented in **Figure G.1** (**Supplementary Materials Section G**). For comparison, we also impute probability density curves using the SEER incidence rates (2022). Given the rare prevalence of *TP53* mutations, one would expect the SEER estimates, which are computed from the general population, to closely match our model-based penetrance estimates for noncarriers.

The major contribution of our approach, however, is the set of covariate-adjusted penetrance estimates for the second primary cancer, which has never been studied in the literature. **Figure 1** shows the corresponding cancer-specific penetrance estimates to the second primary, given a first primary cancer of any type at age 10, from the three individual likelihood models. In contrast, to produce the SEER estimates that match the covariate values, we have to perform the following steps manually. We first select SEER individuals who were diagnosed with first primary cancers of any type at age 10-14 during the period 1975-1989. We then prospectively follow these individuals until 2019 to monitor the occurrences of second primary cancers. The cancer-specific incidence rates at gap times of 5, 10, 15, 20, 25, and 30 years are calculated as the proportions of these individuals who further developed the corresponding cancer type at ages 15-19, 20-24, 25-29, 30-34, 35-39, and 40-44 respectively. In **Figure 1**, our penetrance estimates for individuals with wildtype *TP53* approximates the SEER estimates reasonably well in breast cancer and sarcoma, less well in lung cancer, which has much lower penetrance among SPCs as compared to the other two cancer types. The penetrance estimates of breast cancer and sarcoma are much higher for mutation carriers compared to wildtypes as expected. The penetrance estimates of lung cancer are also higher for mutation carriers, but the difference is much smaller. Although lung cancer belongs to the LFS spectrum, this small difference is not surprising given the very strong competing risks from breast cancer and sarcoma (i.e., mutation carriers are much more likely to develop these two cancer types before lung cancer). While lung cancer has been extensively studied by medical researchers, our study is the first to report age-at-onset penetrance estimates for this cancer type among SPCs. In **Figures G.2–G.3** (**Supplementary Materials Section G**), we provide additional penetrance estimates for two hypothetical patients: one has a first primary cancer at age 20, and the other has a first primary cancer at age 30. These penetrance curves collectively describe how the risk landscapes of patients depend on the ages at diagnosis of their first primary cancers, and demonstrate the utility of our novel statistical framework that formally accounts for such covariates.

**Figure 1:**
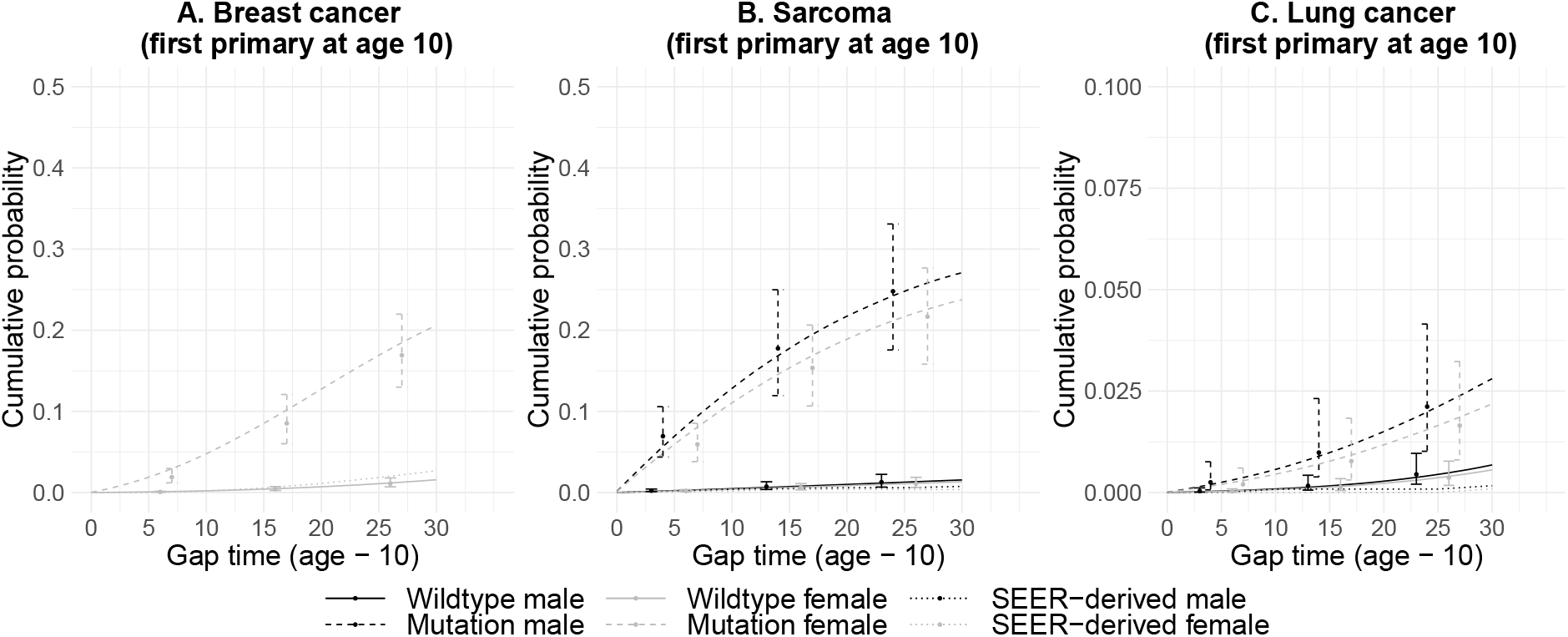
Individual likelihood models: estimates of cancer-specific age-at-onset penetrance to the second primary cancer, given the first primary at age 10. The 95% credible intervals are given at gap times of 5, 15, and 25 years. Cumulative cancer-specific incidence rates from SEER are shown for comparison.

### 3.4 Validation of the penetrance

Let *r_kl_*(*t*|*u*, ***X***(*u*), ***H***), where ***X***(*u*) = {*G, S, D*(*u*)}^*T*^ and ***H*** is the family history, be the risk of developing cancer of type *k* as the *l*th primary by age *t* given the (*l* – 1)th cancer occurrence at age *u*. That is, *r_kl_*(*t*|*u*, ***X***(*u*), ***H***) = *P*(*T_l_* ≤ *t, Z_l_* = *k*|*T*_*l*-1_ = *u*, ***X***(*u*), ***H***). For patients with known *TP53* mutation status, ***H*** can be removed based on the conditional independence assumption. Since we select only family members with measured genotypes, *r_kl_*(*t*|*u*, ***X***(*u*), ***H***) = *P*(*T_l_* ≤ *t, Z_l_* = *k*|*T*_*l*-1_ = *u*, ***X***(*u*)), is equal to the age-at-onset penetrance *q_kl_*(*t*|*u*, ***X***(*u*)). For each cancer model, we evaluate its prediction performance for (1) the first primary, and (2) the second primary for individuals who already had one primary cancer. Receiver Operating Characteristic (ROC) curves for the first evaluation are given in **Supplementary Materials Section H**. The second evaluation is our focus, and the results are shown in **Figure 2**. All models displayed good predictive performances, with areas under the ROC curve (AUCs) being between 0.82 and 0.93, and further presenting small 95% bootstrap confidence intervals.

**Figure 2:**
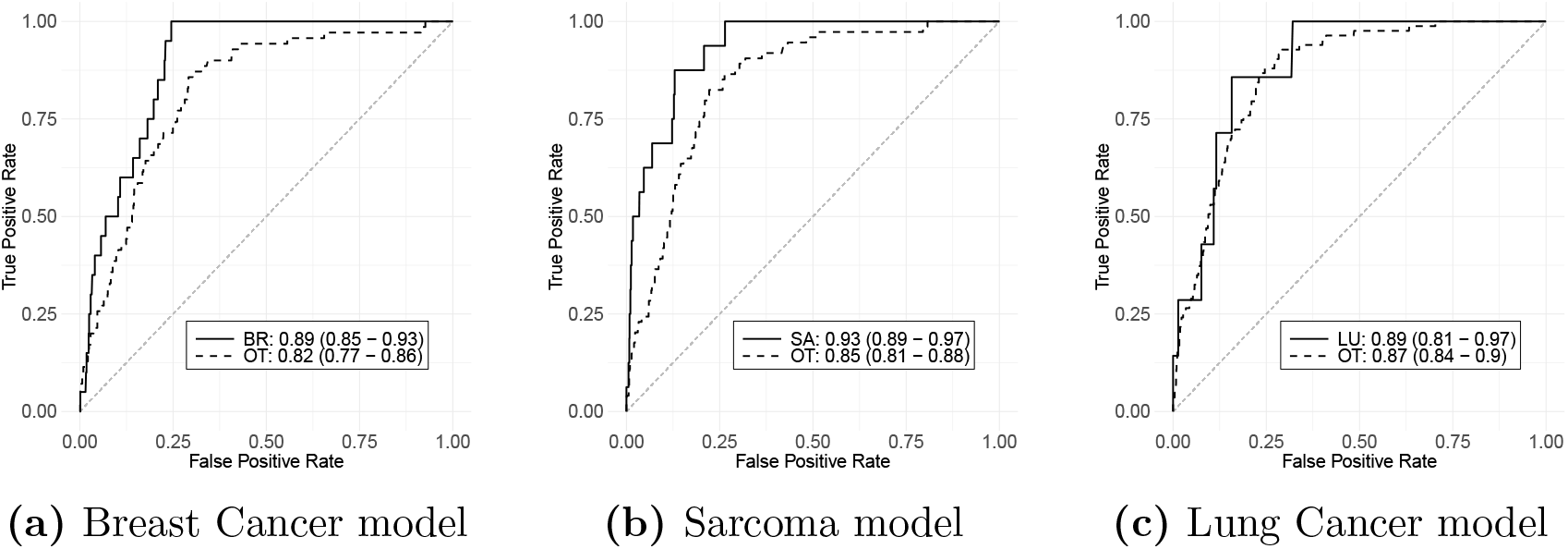
ROC curves, along with the AUCs and their 95% confidence intervals, for cancer-specific prediction of the second primary based on the individual likelihood models. SA = sarcoma, BR = breast cancer, LU = lung cancer, OT = other cancer. Sample size: **(a)** n(BR) = 20, n(OT) = 70; **(b)** n(SA) = 16, n(OT) = 74; **(c)** n(LU) = 7, n(OT) = 83.

## 4 Application of the family-wise likelihood

### 4.1 Model Specifications

Aiming to address the data sparsity issue presented in the application of our individual model to the real dataset, our family-wise model systematically infers missing genotypes and correspondingly includes all family members in the likelihood calculation. Therefore, we fit this model to the full training set across everyone reported. Motivated by both the biologically meaningful outcomes as well as the need for a sufficient sample size for each category, we focus on three groups of cancer diagnosis: (1) sarcoma, (2) breast cancer, and (3) all other cancers combined, which include both LFS-related (i.e., more frequently observed in LFS) and non-LFS malignancies. We include death from other causes as an additional competing risk. For clarity of notations, we set *k* ∈ {*sa, br, ot, de*}, for sarcoma, breast cancer, other cancers, and death respectively. We have ***X***(*t*) = {*G, S, D_sa_*(*t*), *D_br_*(*t*), *D_ot_*(*t*)}^*T*^ and 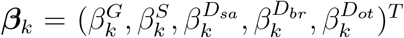 for the regression coefficients in the intensity of cancer type *k*. For sarcoma, other cancers, and death, the intensity functions take the form given by Equation (5). In this dataset, no male patients were diagnosed with breast cancer. Hence, we impose the following restriction on the intensity function of breast cancer: 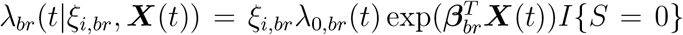. For the ascertainment bias correction, we set *a_sa_* = 10, *a_br_* = 2 and *a_ot_* = 1 as we observe sarcoma to be more indicative of *TP53* mutation.

### 4.2 Model estimation

In addition to the settings described in Section 2.5, we set *M* = 5 for the degree of the Bernstein polynomial to ensure sufficient flexibility while keeping the computational cost reasonable. Using the random walk Metropolis-Hastings-within-Gibbs algorithm, we generate 30,000 posterior samples, of which the first 5,000 are discarded. Overall, the estimation is computationally intensive due to a large number of parameters, with each MCMC iteration taking about 45 seconds on the data when both frailty and ascertainment bias correction is applied. Trace plots of the posterior samples confirm good convergence within 30,000 iterations (**Figures I.1–I.4, Supplementary Material Section I**).

**Table 2** displays all estimates of the regression coefficients from our model. All estimates are significant at 95% significance level, except 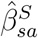, which agrees with a previous study on sarcoma by Amadeo et al. (2020). The 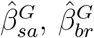, and 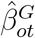 are positive as expected since mutation carriers are much more susceptible to LFS-related cancer types. Parameters of the form 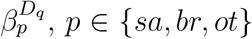 and *q* ∈ {*sa,br, ot*}, are positive with 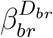 being the sole exception, indicating that patients with a first primary are more likely to develop a second primary later in their lives. A major contributor for this phenomenon is the method of treatment used for the first primary. Radiotherapy and chemotherapy have long been linked to increased risk in cancer survivors (Boice et al., 1985). Bhatia and Sklar (2002) found a strong relationship between radiation-related tumors and the dosage of radiation, and that the second cancers typically develop within the radiation field after a latency period. They also reported some chemotherapeutic agents, such as alkylating agents, that increase the risk of developing second cancer. Sarcoma is particularly likely to occur in patients who were treated with radiotherapy (Patel, 2000), which may explain the largest effect for the onset of second primary cancers is presented by 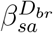. Although data on treatment methods were not collected for our cohort, they can readily be incorporated into our model as an additional covariate in future studies.

**Table 2:** Summary of estimated, *k* ∈ {*sa,br,ot,de*}, based on the last 25,000 posterior samples. The weights *a_sa_* for sarcoma, *a_br_* for breast cancer, and *a_ot_* for other cancers are set to 10, 2, and 1 respectively.

| Event | Parameter | Mean | Median | SD | 2.5% | 97.5% |
| --- | --- | --- | --- | --- | --- | --- |
| Sarcoma | $\beta_{sa}^G$ | 4.063 | 4.063 | 0.223 | 3.645 | 4.494 |
| | $\beta_{sa}^S$ | 0.426 | 0.426 | 0.182 | 0.073 | 0.789 |
| | $\beta_{sa}^{D_{sa}}$ | 1.416 | 1.417 | 0.327 | 0.753 | 2.043 |
| | $\beta_{sa}^{D_{br}}$ | 1.491 | 1.493 | 0.316 | 0.877 | 2.110 |
| | $\beta_{sa}^{D_{ot}}$ | 1.716 | 1.717 | 0.267 | 1.181 | 2.232 |
| Breast | $\beta_{br}^G$ | 3.516 | 3.514 | 0.201 | 3.118 | 3.908 |
| | $\beta_{br}^{D_{sa}}$ | 1.222 | 1.235 | 0.408 | 0.365 | 1.988 |
| | $\beta_{br}^{D_{br}}$ | -0.785 | -0.781 | 0.252 | -1.300 | -0.314 |
| | $\beta_{br}^{D_{ot}}$ | 0.861 | 0.860 | 0.197 | 0.472 | 1.238 |
| Other | $\beta_{ot}^G$ | 2.410 | 2.413 | 0.137 | 2.139 | 2.668 |
| | $\beta_{ot}^S$ | 0.392 | 0.392 | 0.080 | 0.239 | 0.545 |
| | $\beta_{ot}^{D_{sa}}$ | 1.183 | 1.186 | 0.216 | 0.762 | 1.596 |
| | $\beta_{ot}^{D_{br}}$ | 0.641 | 0.644 | 0.169 | 0.295 | 0.964 |
| | $\beta_{ot}^{D_{ot}}$ | 0.783 | 0.783 | 0.113 | 0.557 | 0.993 |
| Death | $\beta_{de}^G$ | 0.841 | 0.836 | 0.132 | 0.578 | 1.102 |
| | $\beta_{de}^S$ | 0.327 | 0.328 | 0.056 | 0.219 | 0.435 |
| | $\beta_{de}^{D_{sa}}$ | 2.309 | 2.314 | 0.157 | 1.987 | 2.614 |
| | $\beta_{de}^{D_{br}}$ | 1.573 | 1.576 | 0.112 | 1.345 | 1.790 |
| | $\beta_{de}^{D_{ot}}$ | 2.139 | 2.138 | 0.061 | 2.016 | 2.259 |

Estimates of ***γ*** and ***ϕ*** are provided in **Tables I.5–I.6** (**Supplementary Material Section I**). To evaluate the sensitivity of our model to the weights *a_k_*, we also report the estimates of the regression coefficients when *a_sa_, a_br_* and *a_ot_* are set to 4, 2 and 1 respectively (**Table I.7, Supplementary Material Section I**). In this case, sarcoma is still more indicative of *TP53* mutation compared to other cancer types, but to a lesser extent. We notice that the estimates do not change much from those in **Table 2**, hence our model is not overly sensitive to the choice of *a_k_*.

### 4.3 Age-at-onset penetrance

We present penetrance estimates for FPC and those for SPC with example onsets of the first primary in sarcoma and other cancers in **Tables J.1–J.5** (**Supplementary Materials Section J**). For comparison, we also show the penetrances that are imputed from the incidence rates provided by SEER (2022). Here we focus on discussing the penetrance estimates for SPC, which have never been reported in the literature. For a female patient with a first primary of breast cancer at age 20, the set of cancer-specific penetrances to the second primary cancer is given by *q*_*k*2_(*t*|20, ***X***), where ***X*** = {*g*, 0,0,1, 0}^*T*^. **Figure 3a** shows the penetrance curves for such a patient with *g* = 0 and *g* =1. The SEER curve was constructed by the following steps. First we selected individuals who were diagnosed with first primary breast cancers at age 20-24 during the period 1975-1989, and prospectively followed them until 2019. We then computed incidence rates at gap times of 5, 10, 15, 20, 25, 30 years as the proportions of individuals who developed second primary cancers of any type during the corresponding age intervals. Similar to **Figure 3b**, we changed the onset of the first primary breast cancer to age 30. With the females who do not carry *TP53* mutations, the SEER estimate approximates closely for those with FPC at age 30 but is much higher than those with FPC at age 20. This highlights an advancement in our model to explicitly adjust for mutation status as a covariate, whereas curves derived from the SEER registry cannot differentiate those with or without *TP53* mutations. For individuals with FPC of breast at age 20, their chances of carrying germline mutations are high. Hence the corresponding SEER curve should approximate more closely to the penetrance estimate for mutation carriers (right panel) than noncarriers (left panel) in **Figure 3a**.

**Figure 3:**
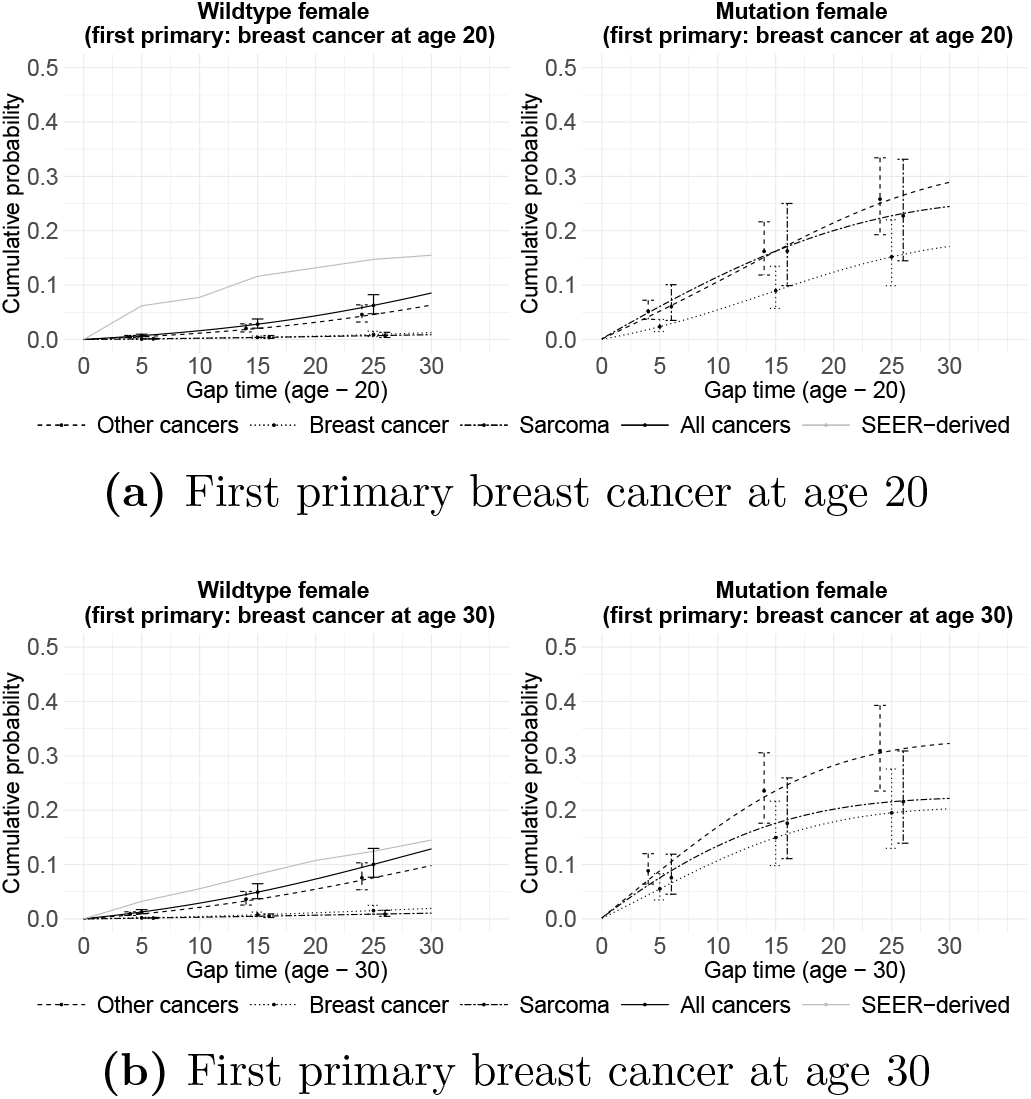
Estimates of cancer-specific age-at-onset penetrance to the second primary cancer, given that the first primary is breast cancer. The 95% credible intervals at gap times of 5, 15, and 25 years are shown. Cumulative incidence rates across all cancer types from SEER are shown for comparison.

### 4.4 Validation of the penetrance

A majority of family members of patients with LFS do not undergo genetic testing for various reasons. Our family-wise likelihood is particularly advantageous in this case because it incorporates all family members’ cancer history into the computation of the time-to-event coefficients. Let ***X***_*G*_(*u*) be the set of covariates without the mutation status (i.e., ***X***_*G*_(*u*) = ***X***(*u*) \ {*G*}). By conditioning on the unobserved *G*, the risk *r_kl_* is given by a weighted average: 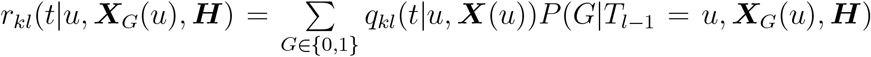 where the mutation probability *P*(*G*|*T*_*l*-1_ = *u*, ***X***_*G*_(*u*), ***H***) is calculated recursively using the peeling algorithm (Elston and Stewart, 1971).

We first evaluate the predictive performance of our joint model in predicting outcomes, such as the *TP53* mutation status, number of primaries, or cancer types within the first primary, which are achievable by previous models (Shin et al. (2019, 2020)). The joint model achieved comparable performances **Supplementary Materials Section K**.

We then move further to predict a unique outcome made available by the joint model, the type of cancer within the second primary. This evaluation is particularly noteworthy as it sets our model apart from others in the literature. Given the limited sample size, we include individuals with mutation probabilities greater than 0.99 as inferred carriers, and smaller than 0.01 as inferred wildtypes. For comparison, we consider a naïve model that assumes independence and identical distributions of the first and second primary. Under this assumption, we can show that the cancer-specific age-at-onset penetrances to the second primary is given by

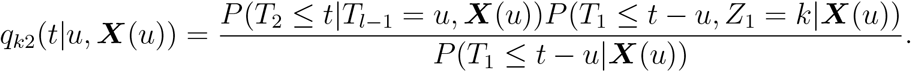

The second term in the numerator is given by a cancer-specific model (Shin et al., 2020), while the other two are given by an MPC model (Shin et al., 2019). We use the penetrance estimates produced by this naive model to perform risk prediction on the validation cohort. **Figure 4** compares the validation results of our model with the naive model. For our model, the AUCs are 0.91, 0.76 and 0.68 for sarcoma, breast cancer, and other cancers, respectively. Furthermore, the ROC curves of the naive model hoover around the diagonal line, with AUC not significantly different from 0.5 for sarcoma and other cancer types. This indicates poor predictive performance and further confirms the importance of accounting for the type and age-at-onset for the first primary when predicting the second primary.

**Figure 4:**
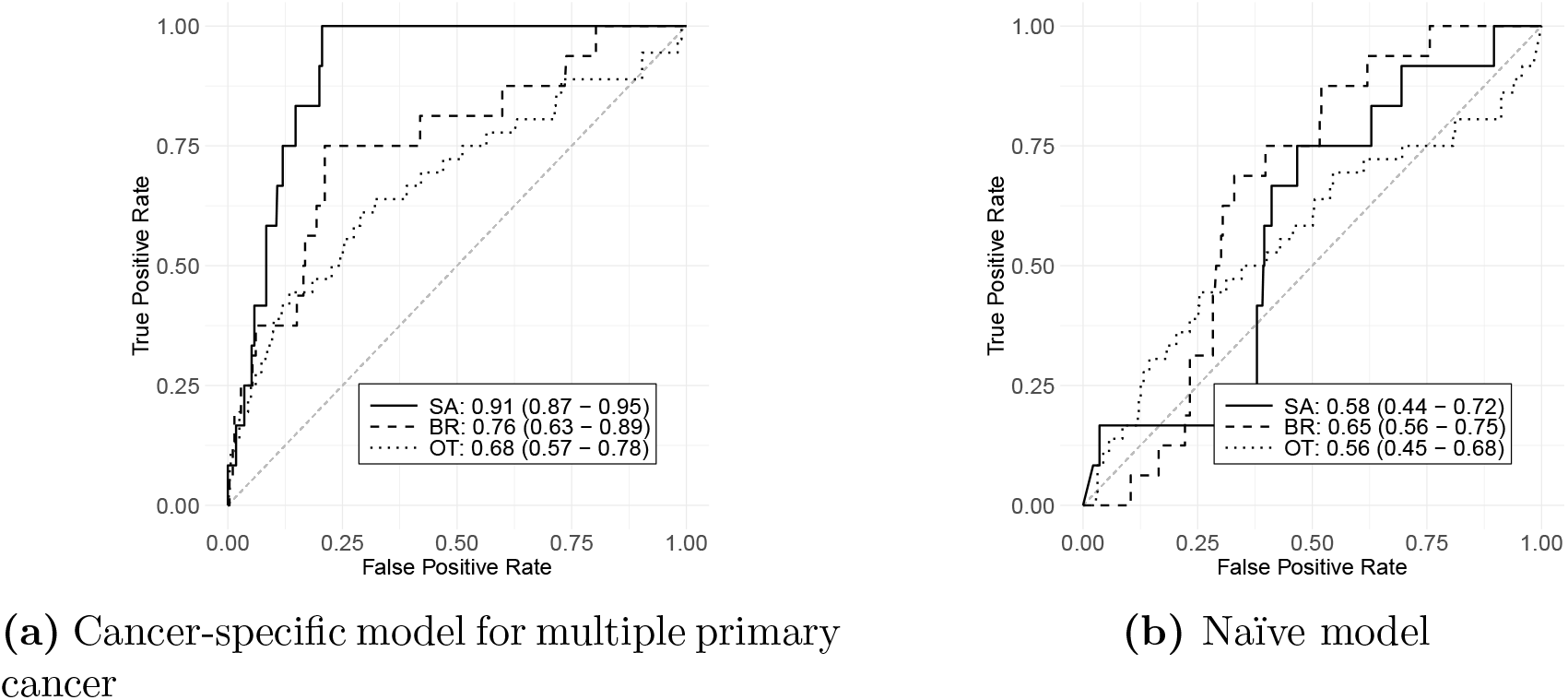
ROC curve, along with the AUCs and their 95% confidence intervals, for cancer-specific prediction of the second primary in the test dataset. SA = sarcoma, BR = breast cancer, OT = other cancer. Sample size: n(SA) = 12, n(BR) = 16, n(OT) = 36.

## 5 Conclusion

With the advancements in oncology, cancer patients are living longer, resulting in a sharp increase in the number of patients diagnosed with multiple primary cancer. Our Bayesian semiparametric framework that jointly models both multiple primary cancers and multiple cancer types represent one of the first attempts to build a quantitative foundation for characterizing cancer risk trajectories among cancer survivors. We provide explicit expressions of both an individual likelihood and a family-wise likelihood, allowing us to evaluate cancer risk trajectories among both independent individuals and family members. The individual likelihood can be used for risk characterization of a general population whose cancer history is captured by registries, while family-wise likelihood can be used for a given inherited disease where family history data are routinely collected. With estimated coefficients by the likelihoods, a suite of penetrance curves can be built, which are cancer-type-specific age-at-onset probabilities for both first and second primary cancers. These probabilities serve as the basis for calculating personalized risk estimates, some of which, for first primary cancer only, are already used in clinical practice to facilitate decision-making (Parmigiani et al., 1998). Our joint model has the potential to move the needle in clinical management by expanding the current practice to the population of cancer survivors.

We have validated the performance of our models using both simulation studies and cross-validation for risk prediction in an extensively collected real dataset. Using the individual likelihood approach, we accurately characterized the risk of second primary lung cancer (SPLC) among cancer survivors in our study population. Our estimated penetrance curves for SPLC is important for the future update of the USPSTF recommendation for lung cancer screening. Using the family-wise likelihood approach, we have obtained a new set of penetrance estimates for genetic and risk counseling with families with LFS. A constant question during the counseling sessions from these families is when my child will develop next cancer and what it will be. Our framework is general and can accommodate any number of cancer types and the number of primary cancers, as well as additional covariates such as treatment information, given additional data availability. Given enriched data with a long-term follow-up in the future, we can apply our model to reveal a complete picture of the risk landscape in cancer patients.

One limitation of our methods is the relatively high computation cost. Training the family-wise model required about three weeks on a high-performance computing cluster, and training individual likelihood models took even longer. To tackle this issue, We will explore moving the computation to Stan, a programming language written in C++ that has been fine-tuned for performance in Bayesian inference.

## Acknowledgments

The authors thank Gang Peng for providing the code to implement the peeling algorithm.

## Supplementary materials

### A. Derivation of individual likelihood

For each individual, the likelihood function is, by definition, given by:

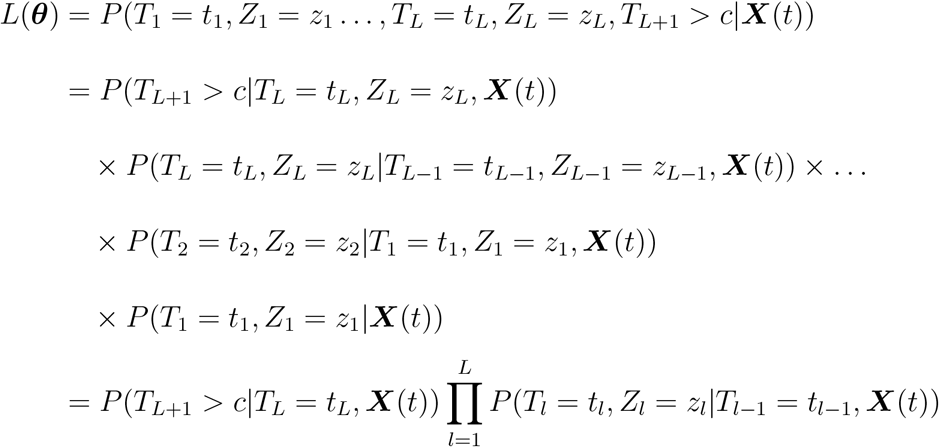

where *T*_0_ = 0 with probability 1. Note that we only need to condition on the most recent event due to the Markov’s property. The last equality assumes that the cancer type of the next occurrence does not depend on the last one. Each probability in the product corresponds to the l-th recurrent event and can be calculated as follows:

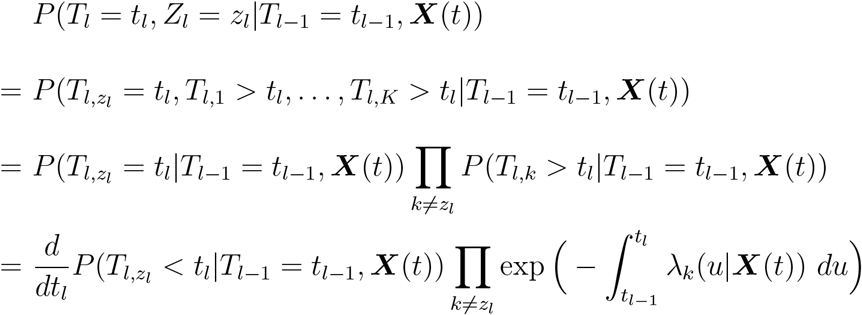

Note that

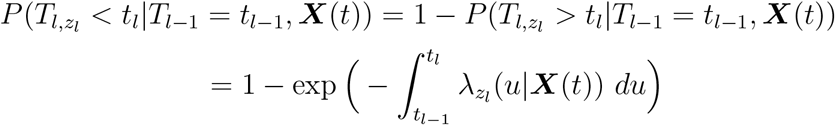

Therefore,

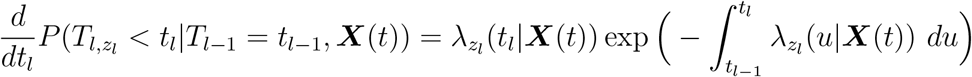

It then follows that

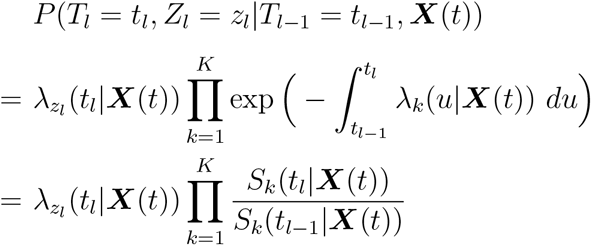

where 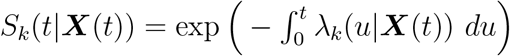.

To complete the likelihood function, we also need to take care of the censored case:

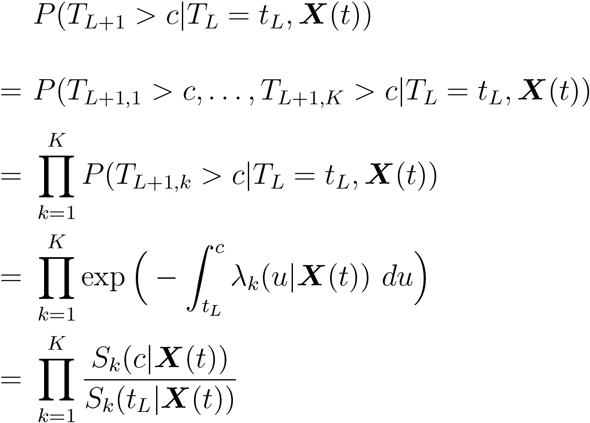

Thus, the full likelihood function for an individual is given by

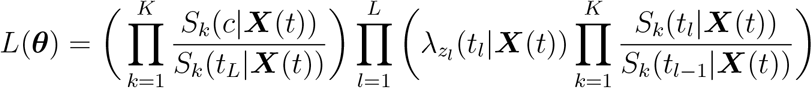

### B. Family-wise likelihood with the peeling algorithm

Following Shin et al (2018), the pedigree structure is partitioned into three disjoint groups: (1) a pivot member *j*, (2) the posterior, which consists of family members that relate to the pivot through his or her spouse or offsprings, and (3) the anterior, which consists of those that relate to the pivot through his or her parents. Let 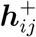 and 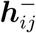 be the cancer histories associated with the posterior and the anterior respectively. Thus, we have partitioned the aggregated family cancer history as 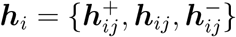.

If *g_ij_* is unobserved, the family-wise likelihood *P*[***h**_i_*|***g***_*i,obs*_ is calculated by conditioning on *g_ij_*

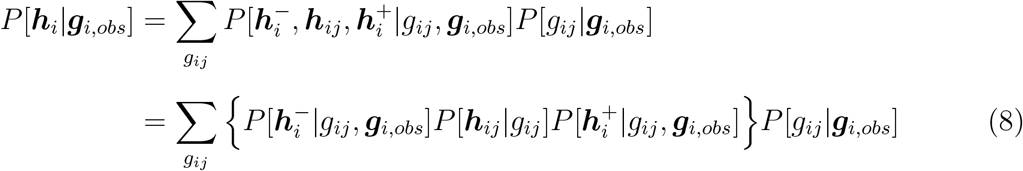

where 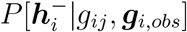 is called the anterior probability and 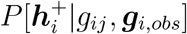 is called the posterior probability.

If *g_ij_* is observed, then it is part of ***g***_*i,obs*_ and the family-wise likelihood is simply

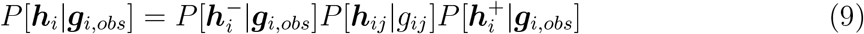

Note that *P*[***h***_*ij*_ |***g***_*ij*_] is the individual likelihood contribution of the pivot member. In either case, the anterior and posterior probabilities can be computed by recursively partitioning 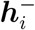 and 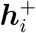 into disjoint subgroups as before. The computation reduces to the evaluation of individual likelihoods, and terms of the type 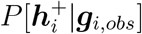, which are easy to compute given that the mode of inheritance is known. Here we assume the Mendelian law of transmission.

### C. Simulation study for individual likelihood with completely observed *G*

To test the individual likelihood, we simulated 20 datasets, each consisting of 10,000 individuals with known genotypes, i.e., mutation carrier (as 1) or noncarrier (as 0) for *TP53*. For simplicity, we assume that there are two cancer types (i.e. *K* = 2). Let 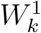 be the gap time to the first occurrence of cancer of type *k*, and 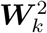 be a vector of size *n* containing the next *N* gap times, where *N* is a large number. It is sufficient to choose *N* = 100. The genotype of the proband is simulated by *G* ~ *Bernoulli*(0.001). Given *G*, 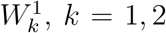, are sampled from exponential distributions with rates

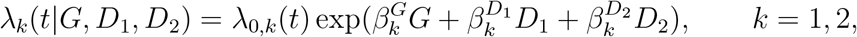

where 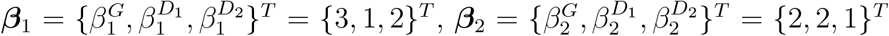, and baseline hazards *λ*_0,1_ (*t*) = *λ*_0,2_(*t*) = 0.001. Because either of the two cancer types can occur as the first primary cancer before the onset of the second primary cancer, we let *D_k_* denote whether cancer type *k* has happened as the first primary, and it follows that *D*_1_ = *D*_2_ = 0 for 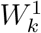. Given 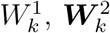, are sampled similarly, except that 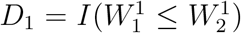 and 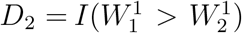. In light of the competing risk framework, we can determine the vector of times at cancer diagnosis, denoted by ***T*** = {*T*_1_, *T*_2_, …, *T*_*N*+1_}^*T*^, and the vector of observed cancer types, denoted by ***Z*** = {*Z*_1_, *Z*_2_, …, *Z*_*N*+1_}^*T*^, by comparing 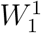 with 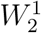, and 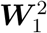 with 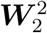 component-wise. The censoring time, denoted by *C*, is randomly generated from Uniform(0, 80). To assess the model performance, we generated 20 independent repetitions.

For parameter estimation, we set *M* = 2 for the degree of the Bernstein polynomials. We then fitted our model to each repetition using MCMC with 10,000 iterations, where the first 5,000 were discarded as burn-in. The trace plots (**Figure C.1**) showed good convergence as well as a good fit for our estimates. Given the true value of a model parameter *β*, we compute the absolute bias of our estimate 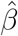

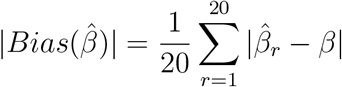

as well as the mean squared error (MSE)

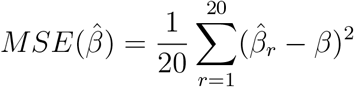

where 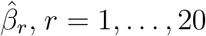, is the estimate of *β* from the *r*-th repetition. The good performance of our parameter estimation on the simulated datasets is further demonstrated by these statistical metrics in **Table C.2**.

**Figure C.1:**
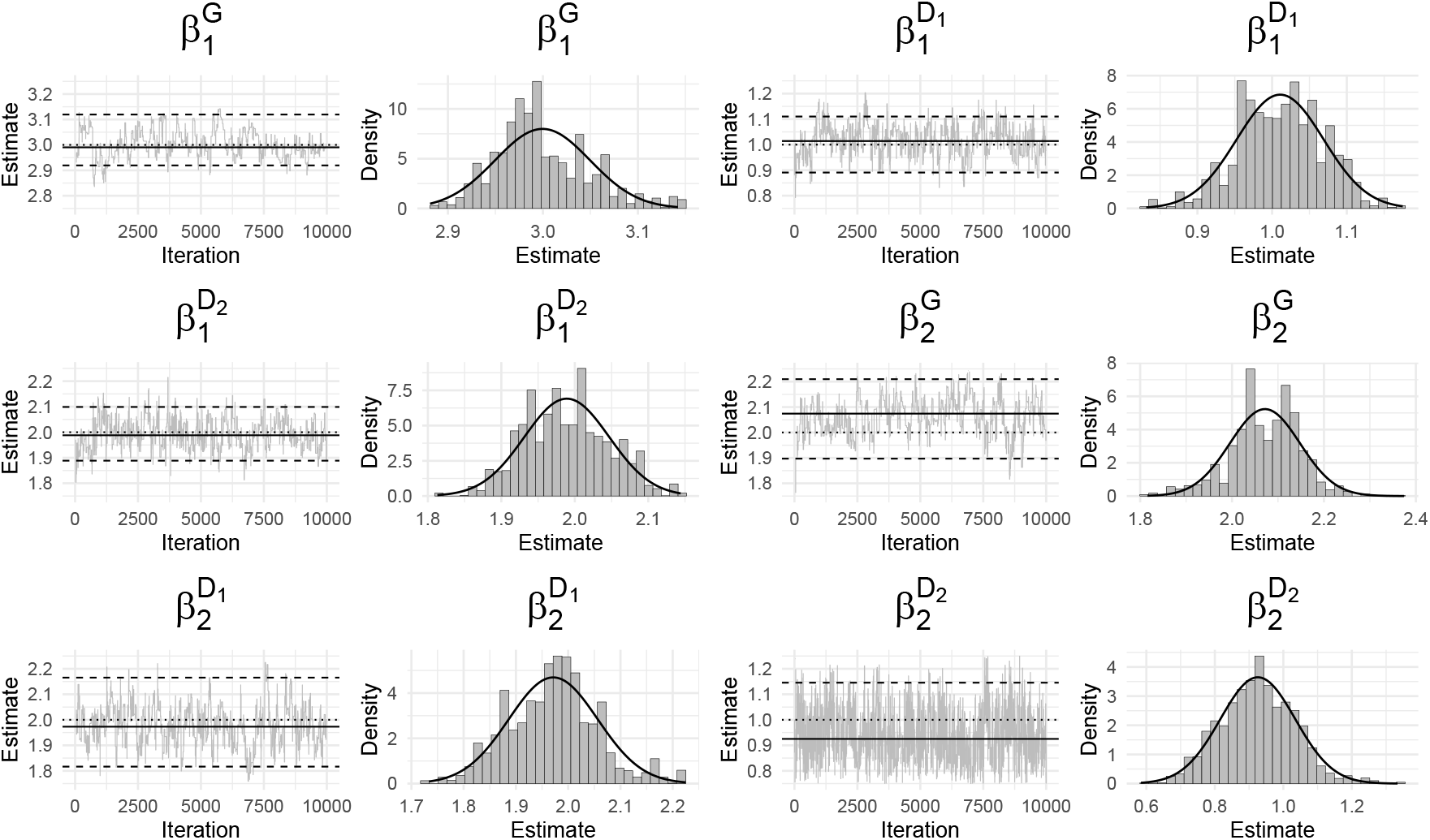
Trace plots of 10,000 posterior samples. Dotted: True parameter; Solid: Estimated parameter; Dashed: 95% credible interval.

**Table C.2:** Absolute biases and mean squared errors (MSE) of parameter estimates in individual likelihood simulation over 20 independent repetitions.

|  | Absolute bias | MSE |
| --- | --- | --- |
| $\beta_1^G$ | 0.064 | 0.005 |
| $\beta_1^{D_1}$ | 0.055 | 0.005 |
| $\beta_1^{D_2}$ | 0.036 | 0.002 |
| $\beta_2^G$ | 0.059 | 0.005 |
| $\beta_2^{D_1}$ | 0.056 | 0.005 |
| $\beta_2^{D_2}$ | 0.097 | 0.012 |
| $\lambda_{0,1}$ | 0.011 | 0.000 |
| $\lambda_{0,2}$ | 0.013 | 0.000 |

### D. Simulation study for family-wise likelihood with partially observed *G*

To test the family-wise likelihood, we simulate 20 datasets, each consisting of 300 families with the same pedigree structure of 30 family members spanning three generations (Figure D.1). For each family, we follow the procedure above to simulate the proband’s cancer history, with the addition of the frailty term

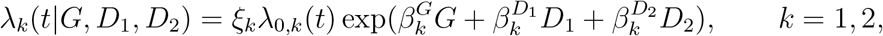

where 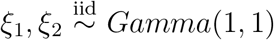. Based on the proband’s data, we generate the genotypes of his/her relatives using the approach described by Shin et al. (2020). Given the genotype, the individual cancer history is independent from other family members, and thus it can be simulated in the same way as the proband. To mimic clinical ascertainment, we use the current clinically used Chompret criteria (the 2015 updated version, as shown in **Table D.2**) (Bougeard et al., 2015) plus a requirement of having at least one *TP53* mutation carrier per family, to ascertain families into an LFS study cohort from the entire simulated population. We repeat the data generation on one new family and its study ascertainment (yes or no) until 300 families are ascertained.

During the parameter estimation, we choose *M* = 2 for the degree of the Bernstein polynomials to keep the computation cost reasonable. For ascertainment bias correction, we set *a*_1_ = *a*_2_ = 1, hence assuming that the ascertainment probability does not depend on the first primary cancer type of the proband. We test a set of models with and without ascertainment bias correction, and with and without the frailty term. In addition, we assess the effectiveness of the peeling algorithm when dealing with missing genotype data, which commonly happens because most family members do not undergo genetic testing. Specifically, we consider (1) no missing genotype, and (2) missing genotype, where a random half of the genotype data are removed. These models are fit to each simulated dataset using MCMC with 30,000 iterations, where the first 5,000 are discarded as burn-in. Finally, all MCMC trace plots (**Figures D.3 – D.6**) show good performance of our parameter estimates. **Table D.7** shows the importance of accounting for ascertainment bias correction and frailty in our model, which generally leads to smaller absolute biases and MSEs. The exceptions are 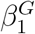 and 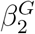, which seem to be best estimated without ascertainment bias correction, with just slightly worse but still good performance after the bias correction.

**Figure D.1.**
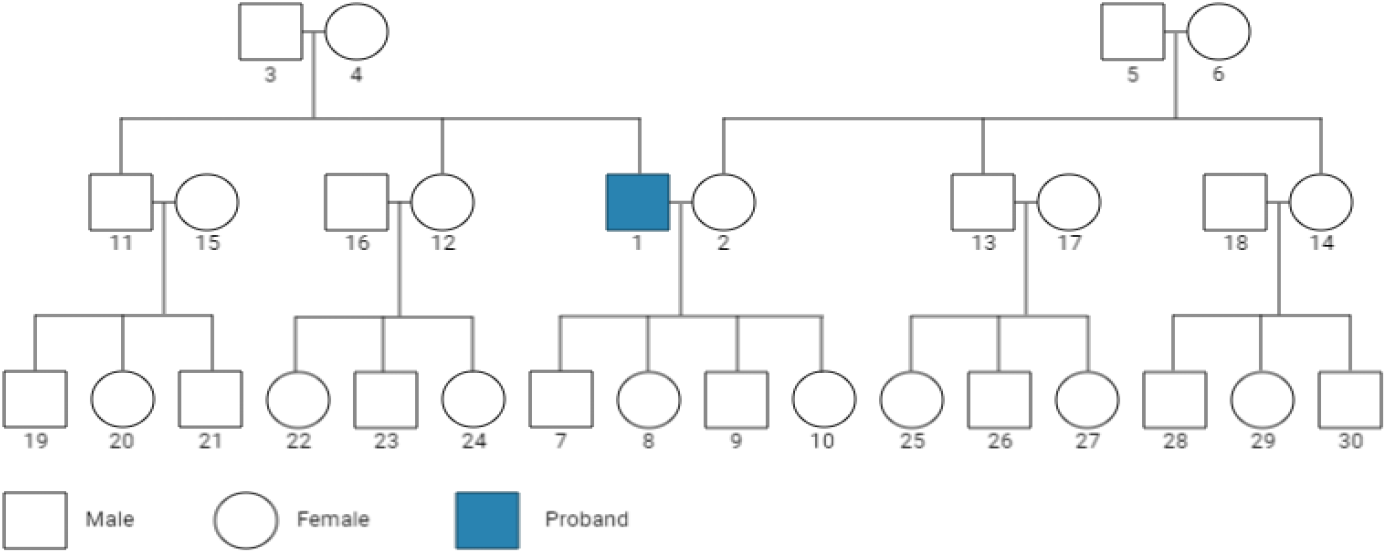
Pedigree structure of a family in a simulated dataset.

**Table D.2:**
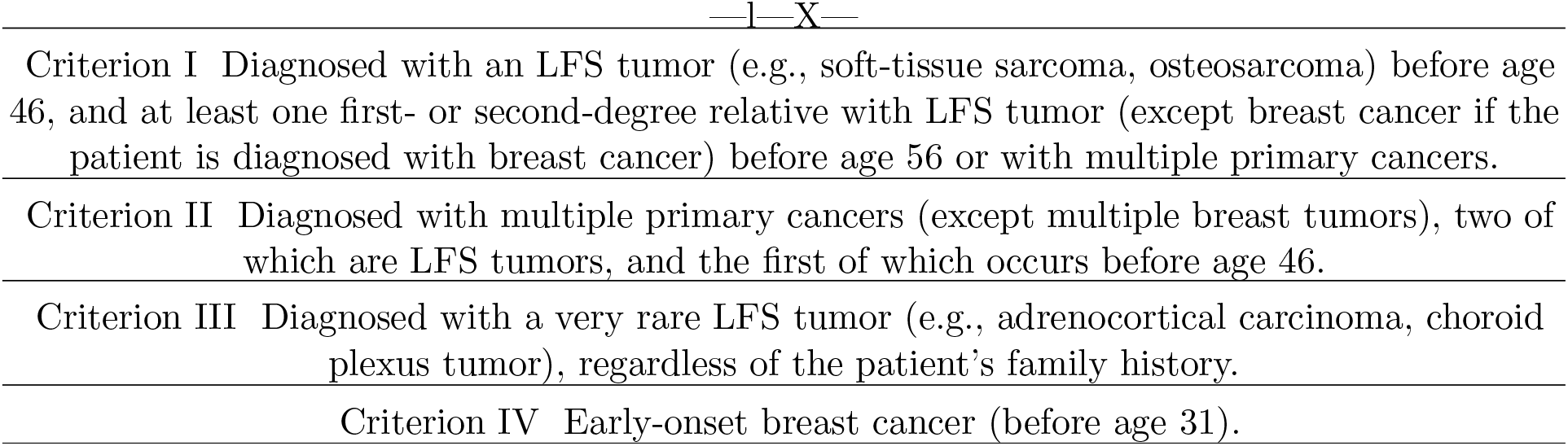
Description of the 2015 revised Chompret criteria to identify patients affected with LFS as those who fall under any single criterion out of the four total. (Bougeard et al., 2015).

**Figure D.3:**
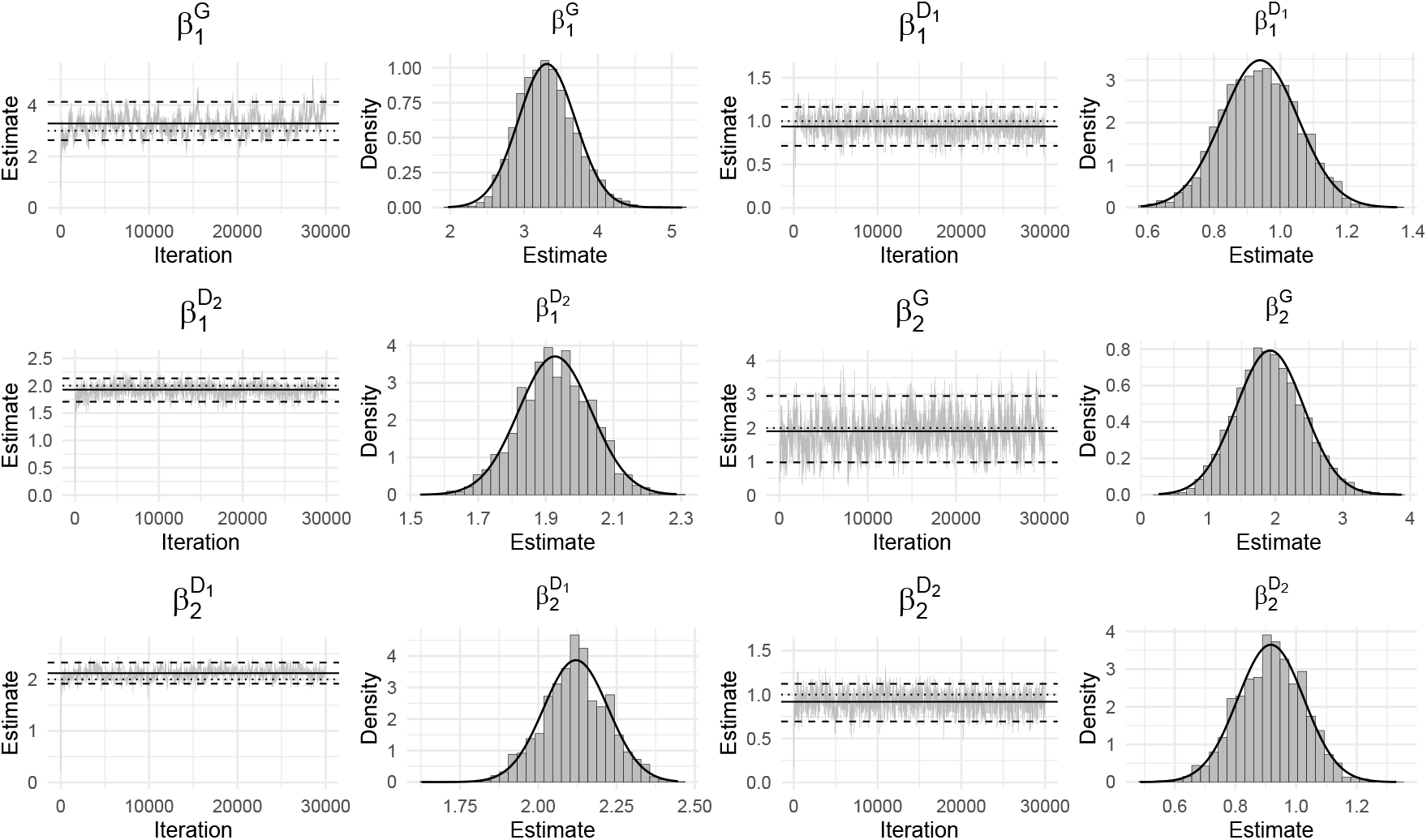
Trace plots of 30,000 posterior samples in the model with both frailty and ascertainment bias correction. Dotted: True parameter; Solid: Estimated parameter; Dashed: 95% credible interval.

**Figure D.4:**
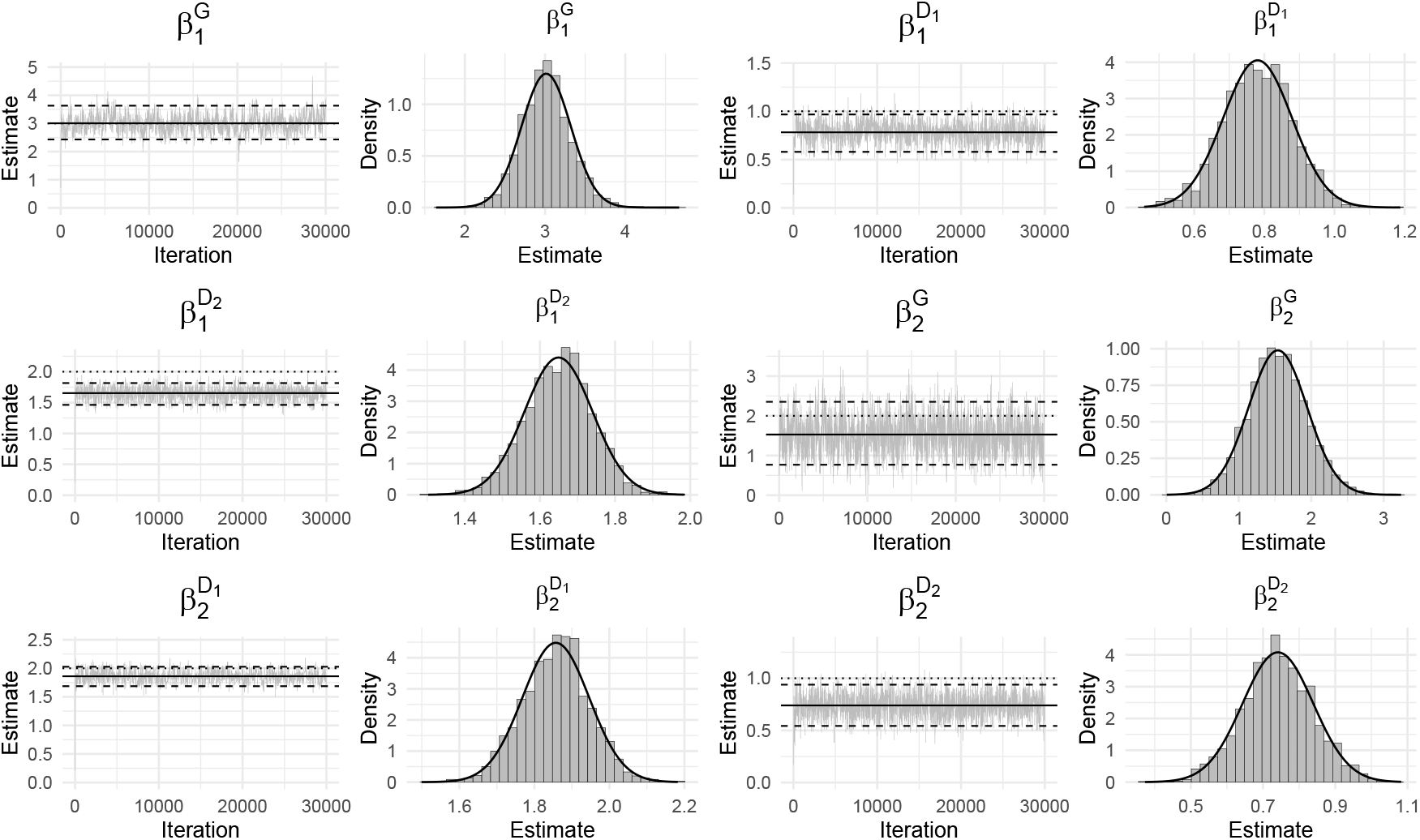
Trace plots of 30,000 posterior samples in the model without ascertainment bias correction. Dotted: True parameter; Solid: Estimated parameter; Dashed: 95% credible interval.

**Figure D.5:**
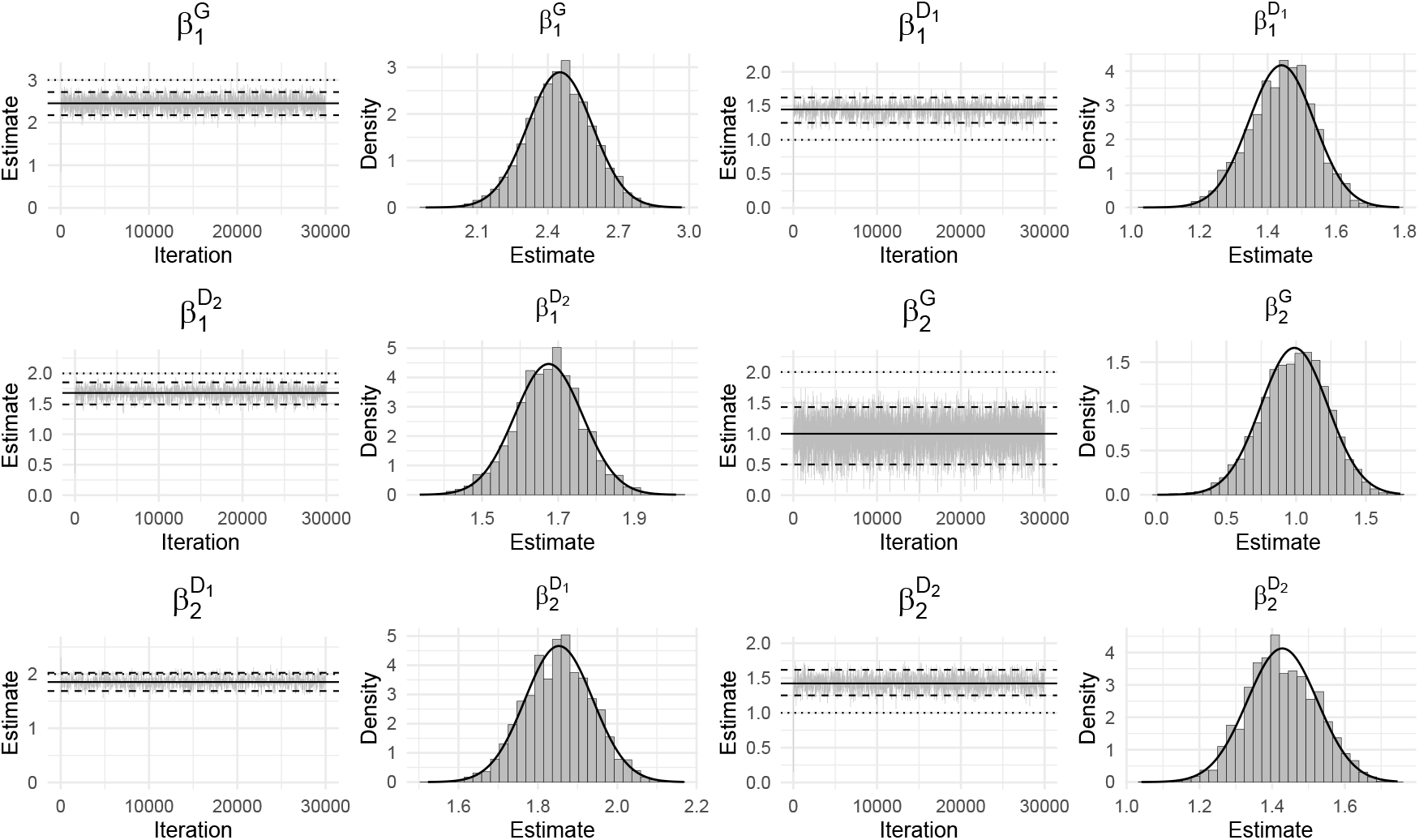
Trace plots of 30,000 posterior samples in the model without frailty. Dotted: True parameter; Solid: Estimated parameter; Dashed: 95% credible interval.

**Figure D.6:**
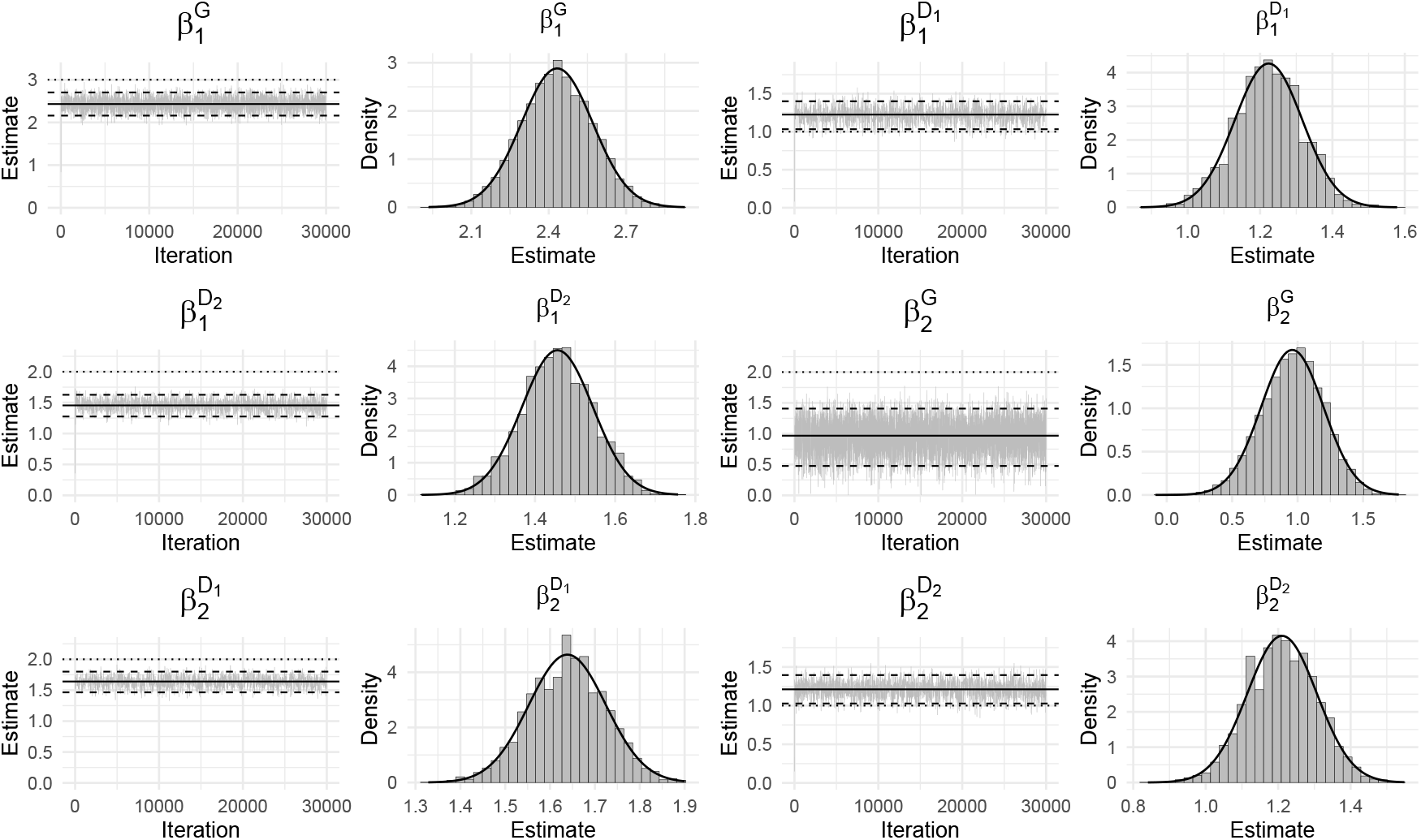
Trace plots of 30,000 posterior samples in the model with neither frailty nor ascertainment bias correction. Dotted: True parameter; Solid: Estimated parameter; Dashed: 95% credible interval.

**Table D.7:**
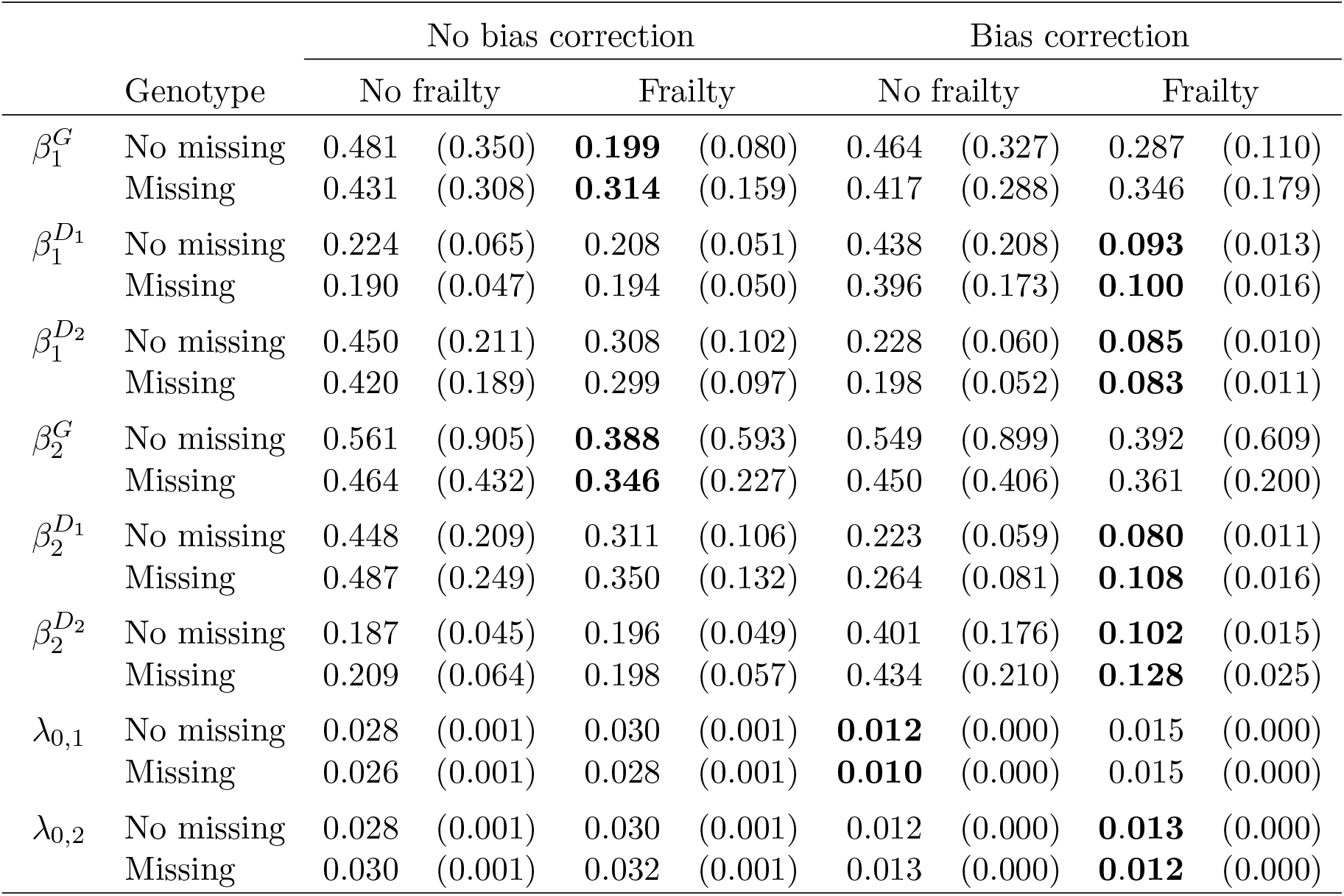
Absolute biases and mean squared errors (in parentheses) of parameter estimates in family-wise likelihood simulation over 20 independent repetitions.

### E. Summary of the MDACC patient cohort data

**Table E.1:** Categorization of family members in the MDACC prospective dataset by gender, mutation status, type of first primary cancer, and type of second primary cancer. BR = breast cancer, SA = sarcoma, OT = other cancers, D = death, C = censored.

|  | First event |  |  |  |  |  | Second event |  |  |  |  |  |
| --- | --- | --- | --- | --- | --- | --- | --- | --- | --- | --- | --- | --- |
|  | BR | SA | OT | D | C | Total | BR | SA | OT | D | C | Total |
| **Male** |  |  |  |  |  |  |  |  |  |  |  |  |
| Wildtype | 0 | 27 | 64 | 3 | 116 | 210 | 0 | 10 | 23 | 22 | 36 | 91 |
| Mutation | 0 | 41 | 48 | 1 | 40 | 130 | 0 | 15 | 24 | 25 | 25 | 89 |
| Unknown | 0 | 70 | 737 | 1131 | 3317 | 5255 | 0 | 6 | 54 | 551 | 196 | 807 |
| Subtotal | 0 | 138 | 849 | 1135 | 3473 | 5595 | 0 | 31 | 101 | 598 | 257 | 987 |
| **Female** |  |  |  |  |  |  |  |  |  |  |  |  |
| Wildtype | 86 | 47 | 94 | 2 | 142 | 371 | 42 | 22 | 50 | 23 | 90 | 227 |
| Mutation | 89 | 36 | 48 | 2 | 30 | 205 | 42 | 24 | 31 | 21 | 55 | 173 |
| Unknown | 350 | 55 | 494 | 881 | 3235 | 5015 | 34 | 8 | 59 | 522 | 276 | 899 |
| Subtotal | 525 | 138 | 636 | 885 | 3407 | 5591 | 118 | 54 | 140 | 566 | 421 | 1299 |
| **Total** | 525 | 276 | 1485 | 2020 | 6880 | 11186 | 118 | 85 | 241 | 1164 | 678 | 2286 |

**Table E.2:** Categorization of family members in the training and validation sets by gender, number of primary cancers and *TP53* mutation status. FPC: first primary cancer. SPC: second primary cancer.

|  | Training set |  |  |  | Validation set |  |  |  |
| --- | --- | --- | --- | --- | --- | --- | --- | --- |
|  | Wildtype | Mutation | Unknown | Total | Wildtype | Mutation | Unknown | Total |
| **Male** |  |  |  |  |  |  |  |  |
| Healthy | 72 | 22 | 2187 | 2281 | 47 | 19 | 2261 | 2327 |
| FPC | 31 | 24 | 388 | 443 | 27 | 26 | 359 | 412 |
| SPC | 21 | 26 | 30 | 77 | 12 | 13 | 30 | 55 |
| Subtotal | 124 | 72 | 2605 | 2801 | 86 | 58 | 2650 | 2794 |
| **Female** |  |  |  |  |  |  |  |  |
| Healthy | 77 | 20 | 2069 | 2166 | 67 | 12 | 2047 | 2126 |
| FPC | 53 | 40 | 388 | 481 | 60 | 36 | 410 | 506 |
| SPC | 55 | 50 | 51 | 156 | 59 | 47 | 50 | 156 |
| Subtotal | 185 | 110 | 2508 | 2803 | 186 | 95 | 2507 | 2788 |
| **Total** | 309 | 182 | 5113 | 5604 | 272 | 153 | 5157 | 5582 |

**Table E.3:** Training data: categorization of family members by gender, number of primary cancers and mutation status, including individuals with mutation probabilities less than 0.01 (inferred wildtype) or greater than 0.9 (inferred mutation). FPC = first primary cancer, SPC = second primary cancer.

|  | Wildtype | Mutation | Inferred wildtype | Inferred mutation | Total |
| --- | --- | --- | --- | --- | --- |
| **Male** |  |  |  |  |  |
| Healthy | 72 | 22 | 1,764 | 10 | 1,868 |
| FPC | 31 | 24 | 182 | 45 | 282 |
| SPC | 21 | 26 | 6 | 9 | 62 |
| Subtotal | 124 | 72 | 1,952 | 64 | 2,365 |
| **Female** |  |  |  |  |  |
| Healthy | 77 | 20 | 1,754 | 2 | 1,853 |
| FPC | 53 | 40 | 230 | 49 | 372 |
| SPC | 55 | 50 | 25 | 10 | 140 |
| Subtotal | 185 | 110 | 2,009 | 61 | 2,212 |
| **Total** | **309** | **182** | **3,961** | **125** | **4,583** |

**Table E.4:** Validation data: categorization of family members by gender, number of primary cancers, and mutation status, including individuals with mutation probabilities less than 0.01 (inferred wildtype) or greater than 0.9 (inferred mutation). FPC = first primary cancer, SPC = second primary cancer.

|  | Wildtype | Mutation | Inferred wildtype | Inferred mutation | Total |
| --- | --- | --- | --- | --- | --- |
| **Male** |  |  |  |  |  |
| Healthy | 47 | 19 | 1,873 | 13 | 1,952 |
| FPC | 27 | 26 | 186 | 64 | 303 |
| SPC | 12 | 13 | 12 | 5 | 42 |
| Subtotal | 86 | 58 | 2,071 | 82 | 2,297 |
| **Female** |  |  |  |  |  |
| Healthy | 67 | 12 | 1,797 | 1 | 1,877 |
| FPC | 60 | 36 | 234 | 59 | 389 |
| SPC | 59 | 47 | 19 | 14 | 139 |
| Subtotal | 186 | 95 | 2,050 | 74 | 2,405 |
| **Total** | **272** | **153** | **4,121** | **156** | **4,708** |

**Table E.5:** Categorization of participants in the SEER Research Plus data by gender, age at diagnosis, type of first primary cancer, and type of second primary cancer. BR = breast cancer, SA = sarcoma, LU = lung cancer.

|  | First primary |  |  | Second primary |  |  |
| --- | --- | --- | --- | --- | --- | --- |
|  | BR | SA | LU | BR | SA | LU |
| **Male** |  |  |  |  |  |  |
| 00 years | 0 | 5 | 4 | 0 | 0 | 0 |
| 01-04 years | 0 | 38 | 10 | 0 | 3 | 0 |
| 05-09 years | 0 | 194 | 3 | 0 | 11 | 1 |
| 10-14 years | 0 | 489 | 19 | 0 | 22 | 1 |
| 15-19 years | 0 | 669 | 48 | 0 | 33 | 1 |
| 20-24 years | 2 | 368 | 98 | 0 | 25 | 3 |
| 25-29 years | 7 | 279 | 185 | 0 | 24 | 6 |
| 30-34 years | 26 | 234 | 526 | 0 | 42 | 27 |
| 35-39 years | 52 | 246 | 1423 | 6 | 44 | 54 |
| 40-44 years | 114 | 237 | 3694 | 1 | 55 | 148 |
| 45-49 years | 219 | 233 | 8557 | 11 | 88 | 433 |
| 50-54 years | 293 | 264 | 16871 | 23 | 98 | 1150 |
| 55-59 years | 456 | 260 | 27933 | 42 | 186 | 2601 |
| 60-64 years | 536 | 258 | 38485 | 61 | 212 | 4896 |
| 65-69 years | 556 | 234 | 45313 | 115 | 284 | 7756 |
| 70-74 years | 483 | 194 | 43109 | 124 | 361 | 9501 |
| 75-79 years | 373 | 142 | 34986 | 149 | 368 | 9431 |
| **Female** |  |  |  |  |  |  |
| 00 years | 0 | 13 | 4 | 0 | 1 | 0 |
| 01-04 years | 1 | 43 | 9 | 0 | 2 | 0 |
| 05-09 years | 1 | 178 | 3 | 0 | 10 | 0 |
| 10-14 years | 8 | 421 | 10 | 1 | 20 | 0 |
| 15-19 years | 42 | 353 | 53 | 4 | 19 | 0 |
| 20-24 years | 479 | 233 | 101 | 18 | 26 | 3 |
| 25-29 years | 3093 | 204 | 190 | 168 | 38 | 11 |
| 30-34 years | 9676 | 187 | 472 | 562 | 46 | 29 |
| 35-39 years | 21593 | 176 | 1207 | 1493 | 55 | 116 |
| 40-44 years | 39236 | 180 | 3078 | 3169 | 85 | 341 |
| 45-49 years | 56739 | 226 | 6496 | 5598 | 108 | 799 |
| 50-54 years | 62683 | 189 | 11932 | 7424 | 143 | 1678 |
| 55-59 years | 65079 | 179 | 18378 | 9297 | 180 | 3021 |
| 60-64 years | 67783 | 215 | 24531 | 11577 | 250 | 4797 |
| 65-69 years | 65899 | 173 | 29144 | 13254 | 250 | 6467 |
| 70-74 years | 56808 | 153 | 30288 | 12898 | 272 | 7462 |
| 75-79 years | 45805 | 109 | 26357 | 11609 | 302 | 6927 |

### F. Estimation of Model Parameters in the Individual Likelihood Models

**Figure F.1:**
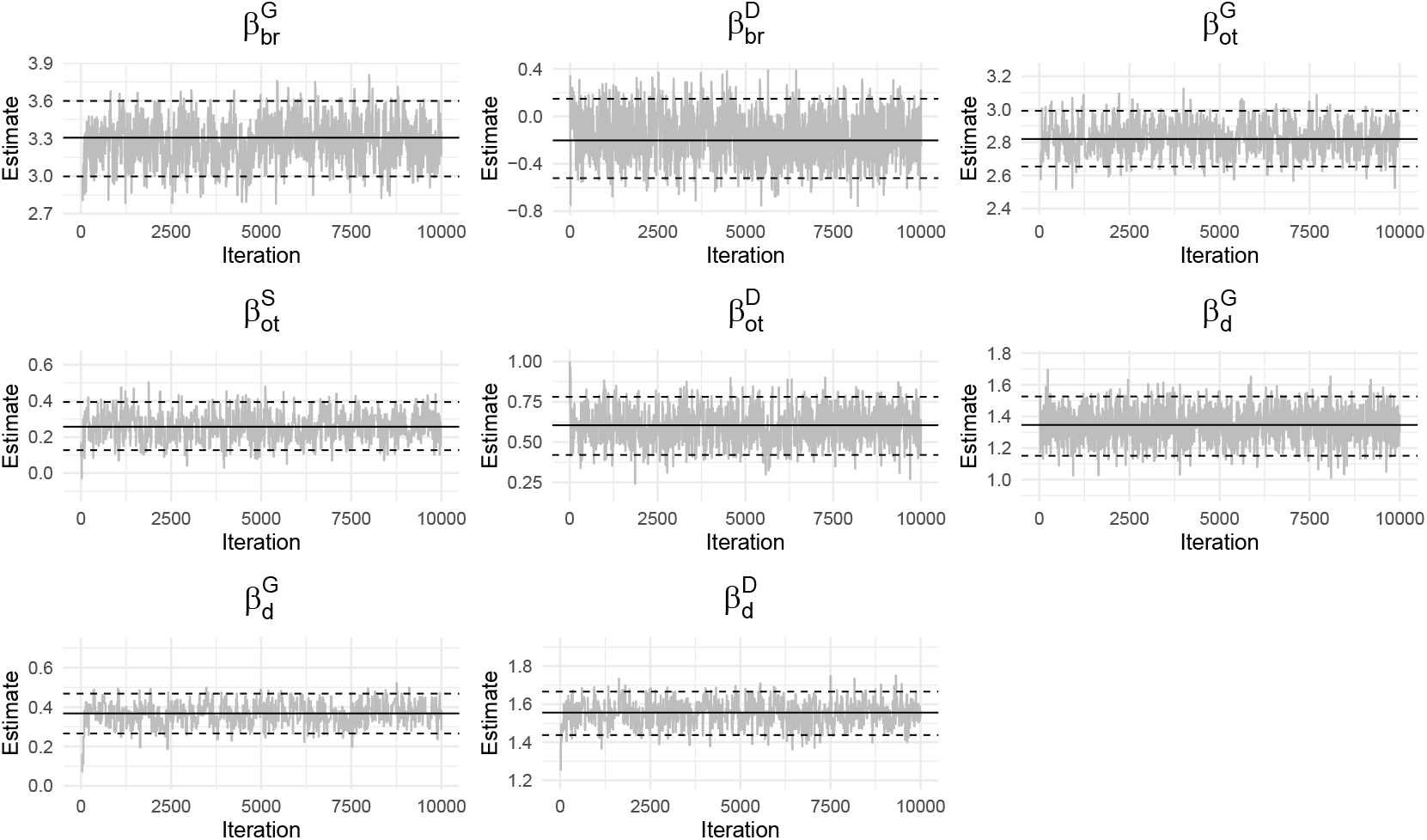
Trace plots of 10,000 posterior samples of the regression coefficients ***β*** in the Breast Cancer model. Solid: Estimated parameter; Dashed: 95% credible interval.

**Figure F.2:**
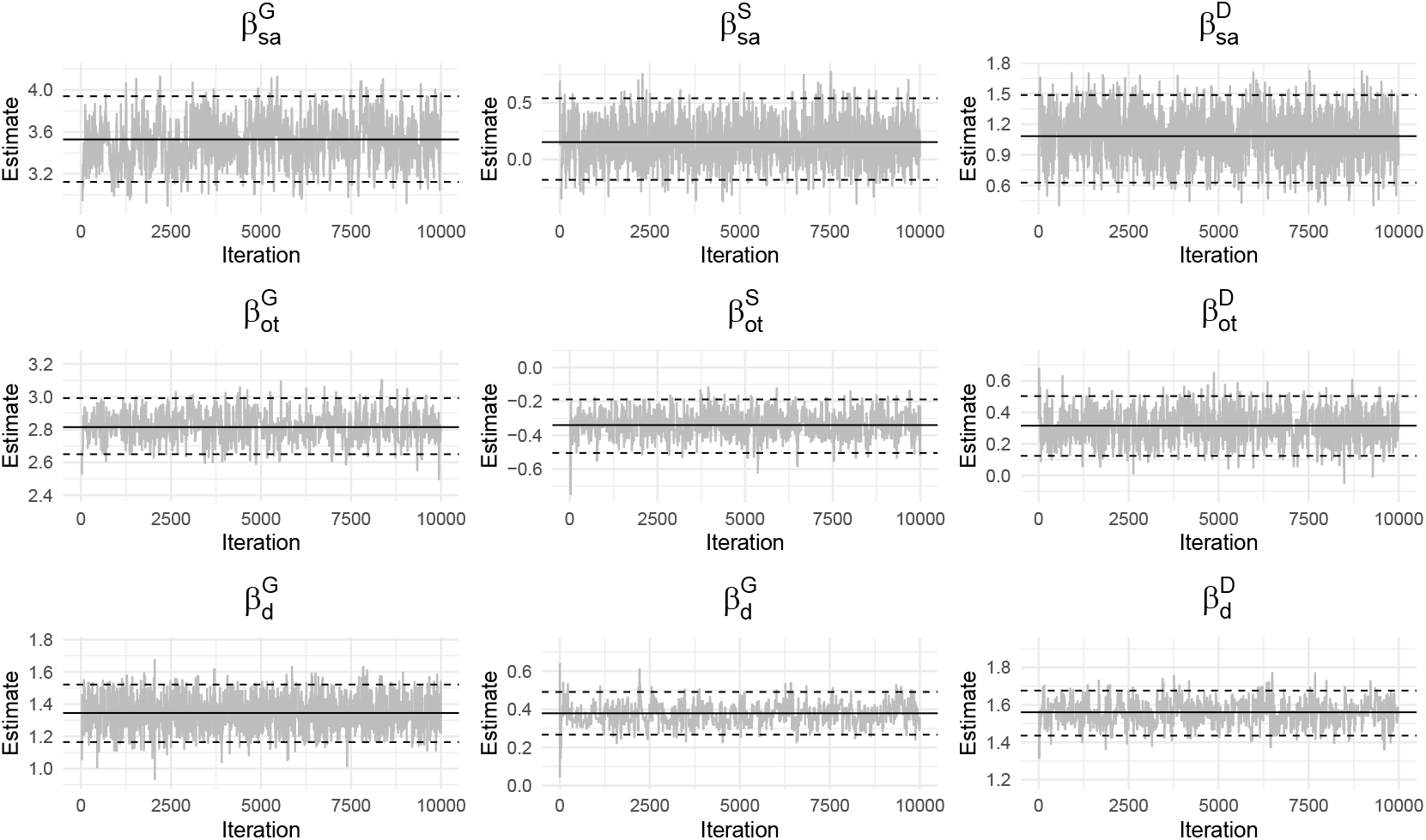
Trace plots of 10,000 posterior samples of the regression coefficients ***β*** in the Sarcoma model. Solid: Estimated parameter; Dashed: 95% credible interval.

**Figure F.3:**
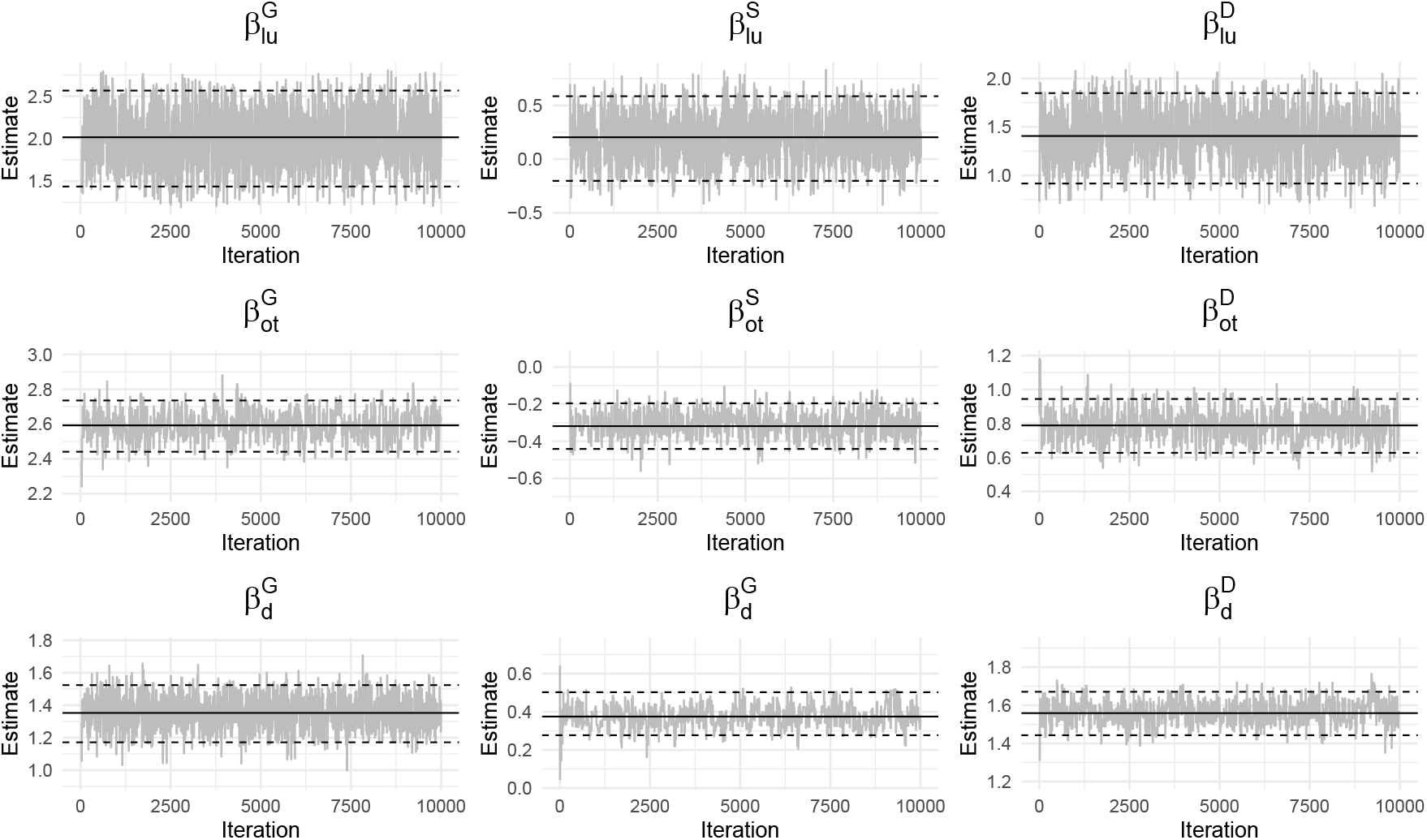
Trace plots of 10,000 posterior samples of the regression coefficients ***β*** in the Lung Cancer model. Solid: Estimated parameter; Dashed: 95% credible interval.

**Table F.4:** Estimated ***β*** in the individual likelihood models based on the last 5,000 posterior samples.

| Model | Event | Parameter | Mean | Median | SD | 2.5% | 97.5% |
| --- | --- | --- | --- | --- | --- | --- | --- |
| Breast Cancer | Breast | $\beta_{br}^G$ | 3.302 | 3.305 | 0.158 | 2.997 | 3.598 |
| | | $\beta_{br}^D$ | -0.200 | -0.203 | 0.170 | -0.522 | 0.148 |
| | Other | $\beta_{ot}^G$ | 2.821 | 2.821 | 0.090 | 2.654 | 2.991 |
| | | $\beta_{ot}^S$ | 0.260 | 0.257 | 0.069 | 0.127 | 0.394 |
| | | $\beta_{ot}^D$ | 0.604 | 0.605 | 0.098 | 0.420 | 0.781 |
| | Death | $\beta_{de}^G$ | 1.346 | 1.346 | 0.093 | 1.151 | 1.526 |
| | | $\beta_{de}^S$ | 0.367 | 0.368 | 0.051 | 0.266 | 0.468 |
| | | $\beta_{de}^D$ | 1.553 | 1.556 | 0.061 | 1.438 | 1.666 |
| Sarcoma | Sarcoma | $\beta_{sa}^G$ | 3.526 | 3.527 | 0.206 | 3.123 | 3.939 |
| | | $\beta_{sa}^S$ | 0.162 | 0.153 | 0.174 | -0.180 | 0.540 |
| | | $\beta_{sa}^D$ | 1.080 | 1.083 | 0.221 | 0.627 | 1.486 |
| | Other | $\beta_{ot}^G$ | 2.816 | 2.815 | 0.085 | 2.650 | 2.991 |
| | | $\beta_{ot}^S$ | -0.341 | -0.340 | 0.074 | -0.505 | -0.189 |
| | | $\beta_{ot}^D$ | 0.314 | 0.316 | 0.096 | 0.124 | 0.503 |
| | Death | $\beta_{de}^G$ | 1.344 | 1.345 | 0.094 | 1.165 | 1.521 |
| | | $\beta_{de}^S$ | 0.379 | 0.379 | 0.053 | 0.267 | 0.491 |
| | | $\beta_{de}^D$ | 1.557 | 1.561 | 0.062 | 1.435 | 1.676 |
| Lung Cancer | Lung | $\beta_{lu}^G$ | 2.017 | 2.017 | 0.296 | 1.436 | 2.567 |
| | | $\beta_{lu}^S$ | 0.198 | 0.204 | 0.198 | -0.202 | 0.585 |
| | | $\beta_{lu}^D$ | 1.409 | 1.407 | 0.239 | 0.916 | 1.850 |
| | Other | $\beta_{ot}^G$ | 2.593 | 2.593 | 0.076 | 2.441 | 2.736 |
| | | $\beta_{ot}^S$ | -0.316 | -0.319 | 0.065 | -0.441 | -0.196 |
| | | $\beta_{ot}^D$ | 0.788 | 0.789 | 0.082 | 0.627 | 0.945 |
| | Death | $\beta_{de}^G$ | 1.349 | 1.352 | 0.091 | 0.171 | 1.524 |
| | | $\beta_{de}^S$ | 0.381 | 0.375 | 0.057 | 0.277 | 0.503 |
| | | $\beta_{de}^D$ | 1.559 | 1.559 | 0.060 | 1.443 | 1.671 |

### G. Age-at-onset Penetrance from the Individual Likelihood Models

**Figure G.1:**
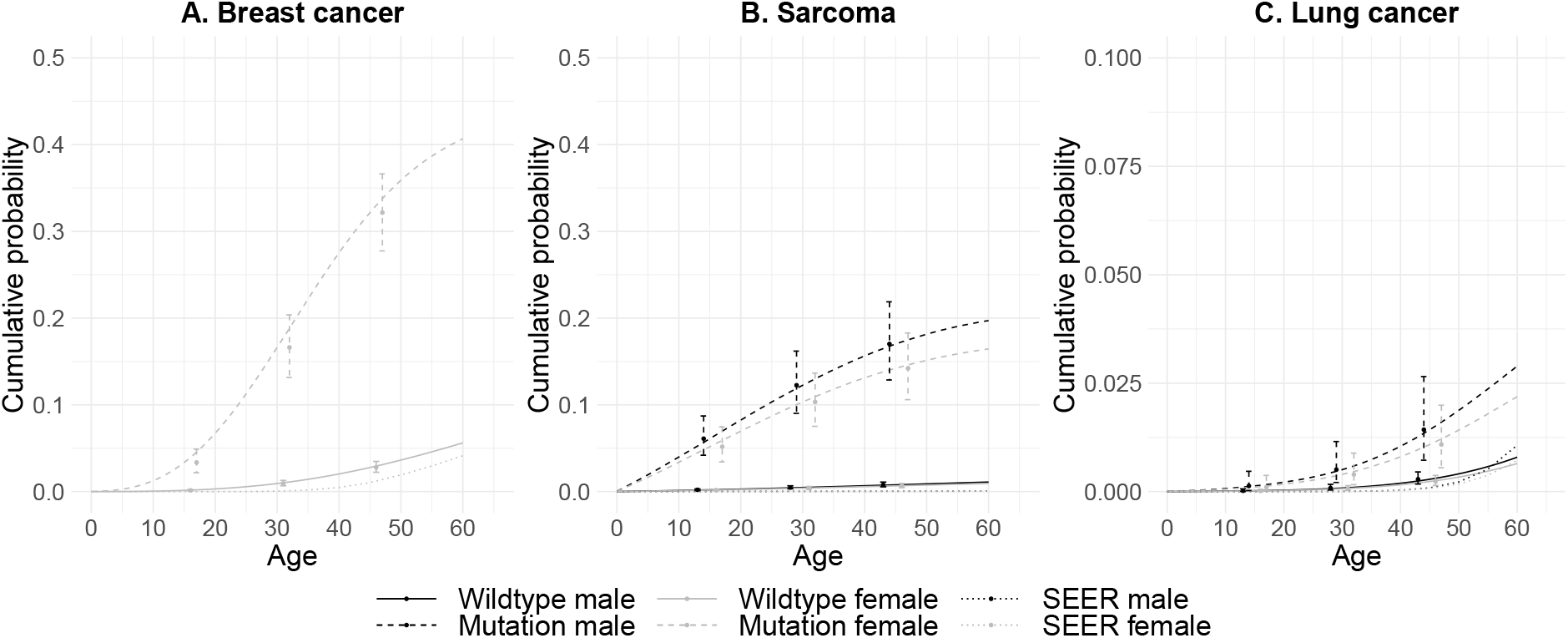
Estimates of cancer-specific age-at-onset penetrance to the first primary cancer for male and female with and without TP53 mutation based on the individual likelihood models, and the corresponding 95% credible intervals at ages 15, 30, and 45. Cancer-specific incidence rates from SEER are shown for comparison.

**Figure G.2:**
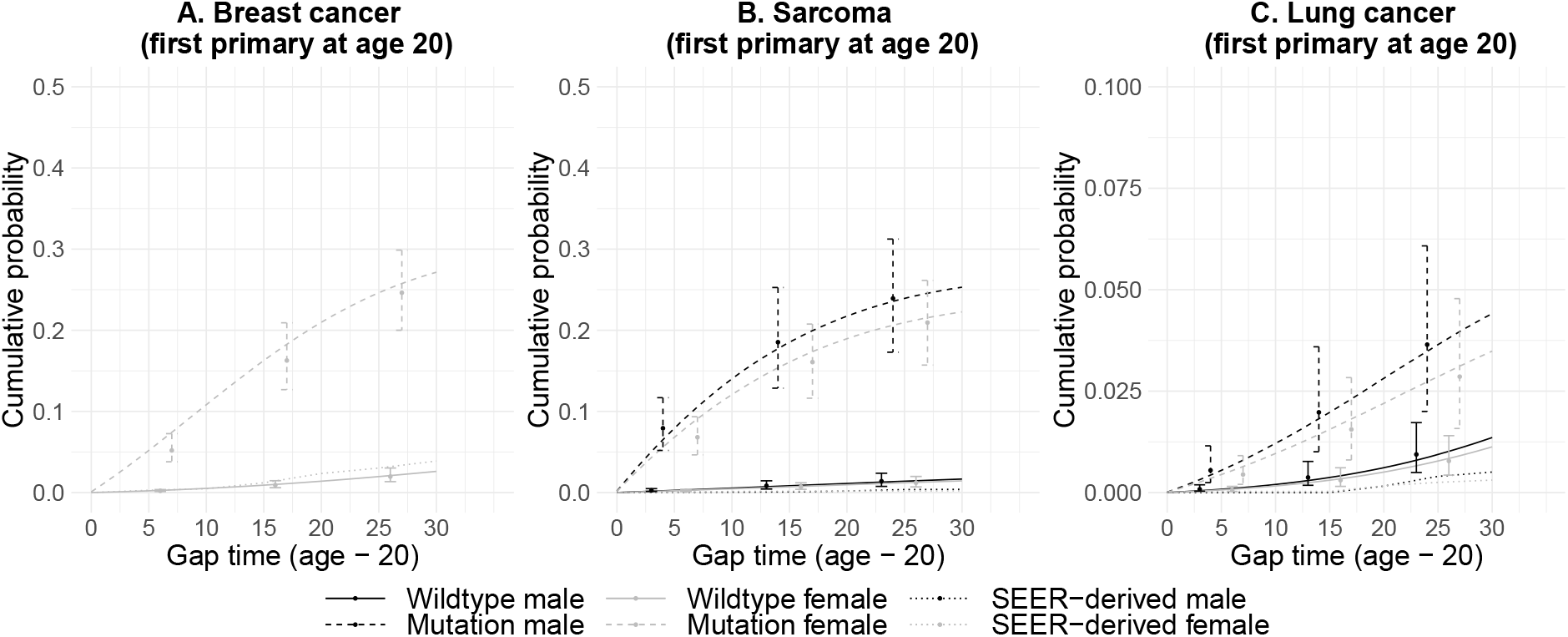
Estimates of cancer-specific age-at-onset penetrance to the second primary cancer, given the first primary at age 20, for male and female with and without *TP53* mutation based on the individual likelihood models, and the corresponding 95% credible intervals at gap times 5, 15, and 25. Cancer-specific incidence rates from SEER are shown for comparison.

**Figure G.3:**
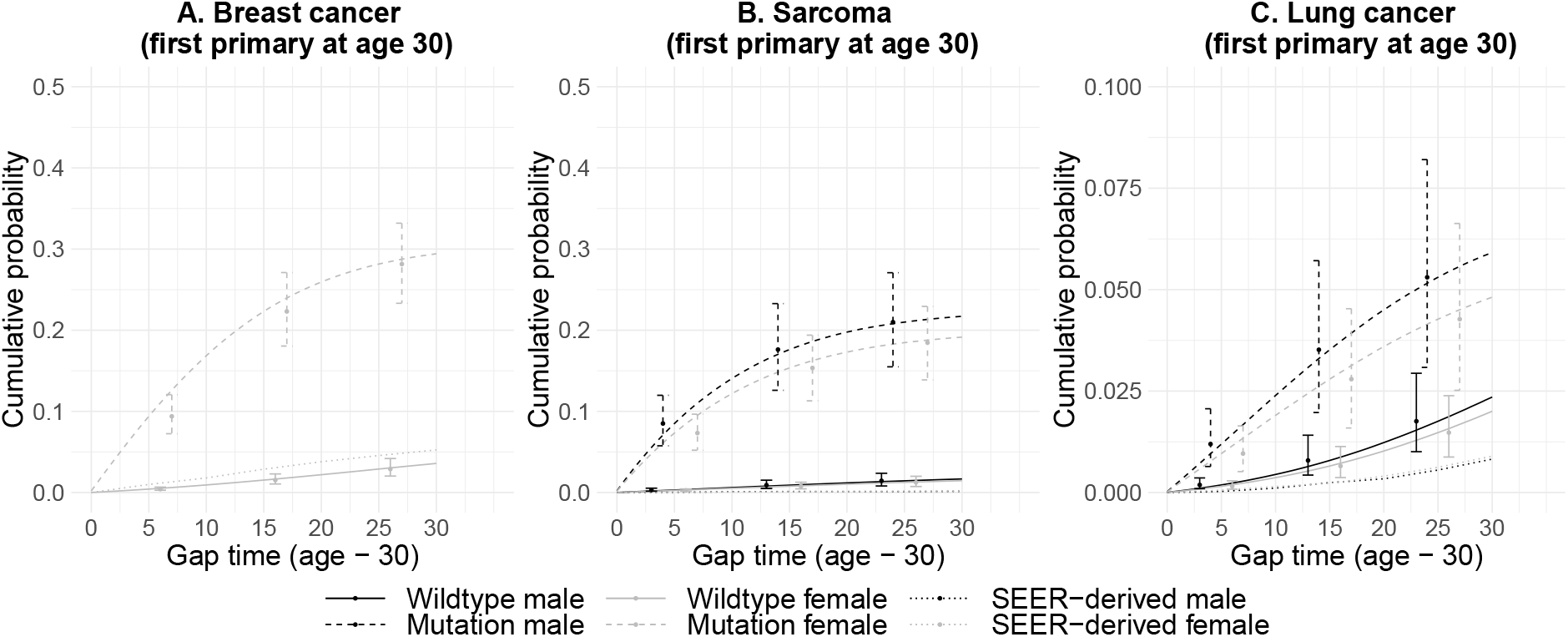
Estimates of cancer-specific age-at-onset penetrance to the second primary cancer, given the first primary at age 30, for male and female with and without *TP53* mutation based on the individual likelihood models, and the corresponding 95% credible intervals at gap times 5, 15, and 25. Cancer-specific incidence rates from SEER are shown for comparison.

### H. Analyses Results from the Validation Study of the Individual Likelihood Models

**Figure H.1:**
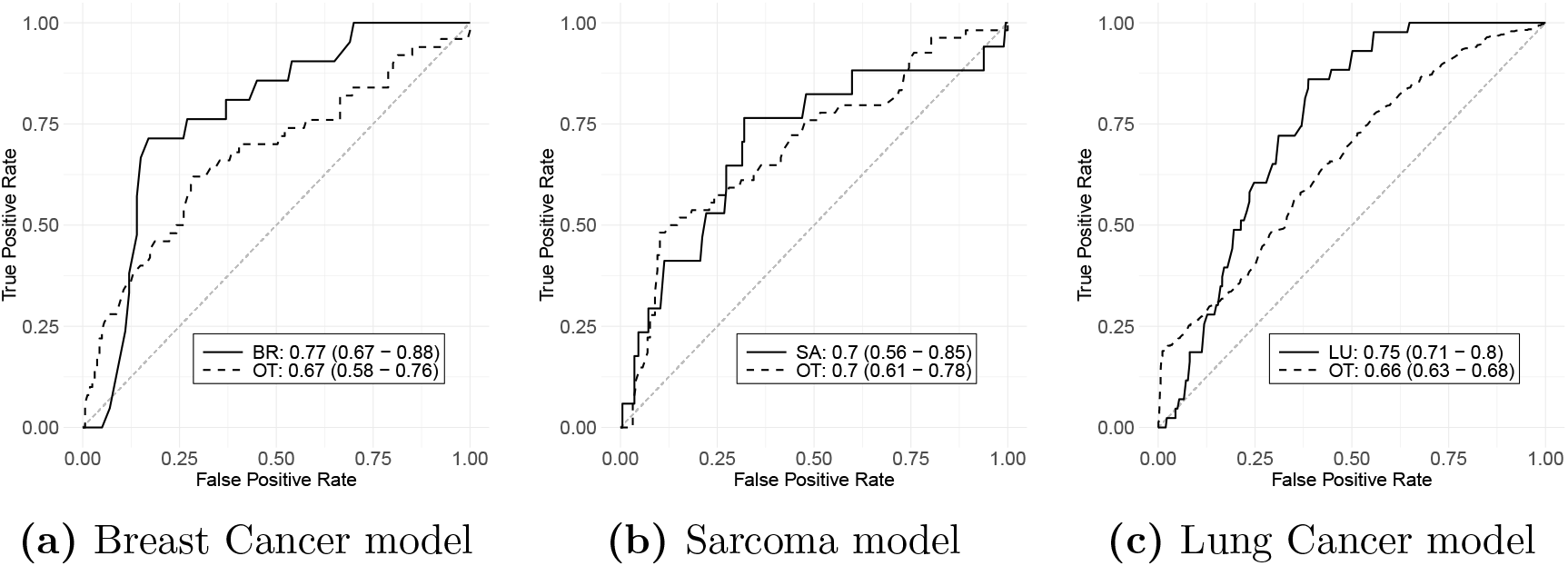
ROC curves, and the 95% bootstrap confidence intervals of the AUCs, for cancer-specific prediction of the first primary cancer in the derived validation dataset. Inferred mutation and non-carriers are included in **(c)** only. BR = breast cancer, SA = sarcoma, LU = lung cancer, OT = other cancer. Sample size: **(a)** n(BR) = 21, n(OT) = 50; **(b)** n(SA) = 17, n(OT) = 54; **(c)** n(LU) = 44, n(OT) = 618

### I. Estimation of Model Parameters in the Family-wise Likelihood Model

**Figure I.1:**
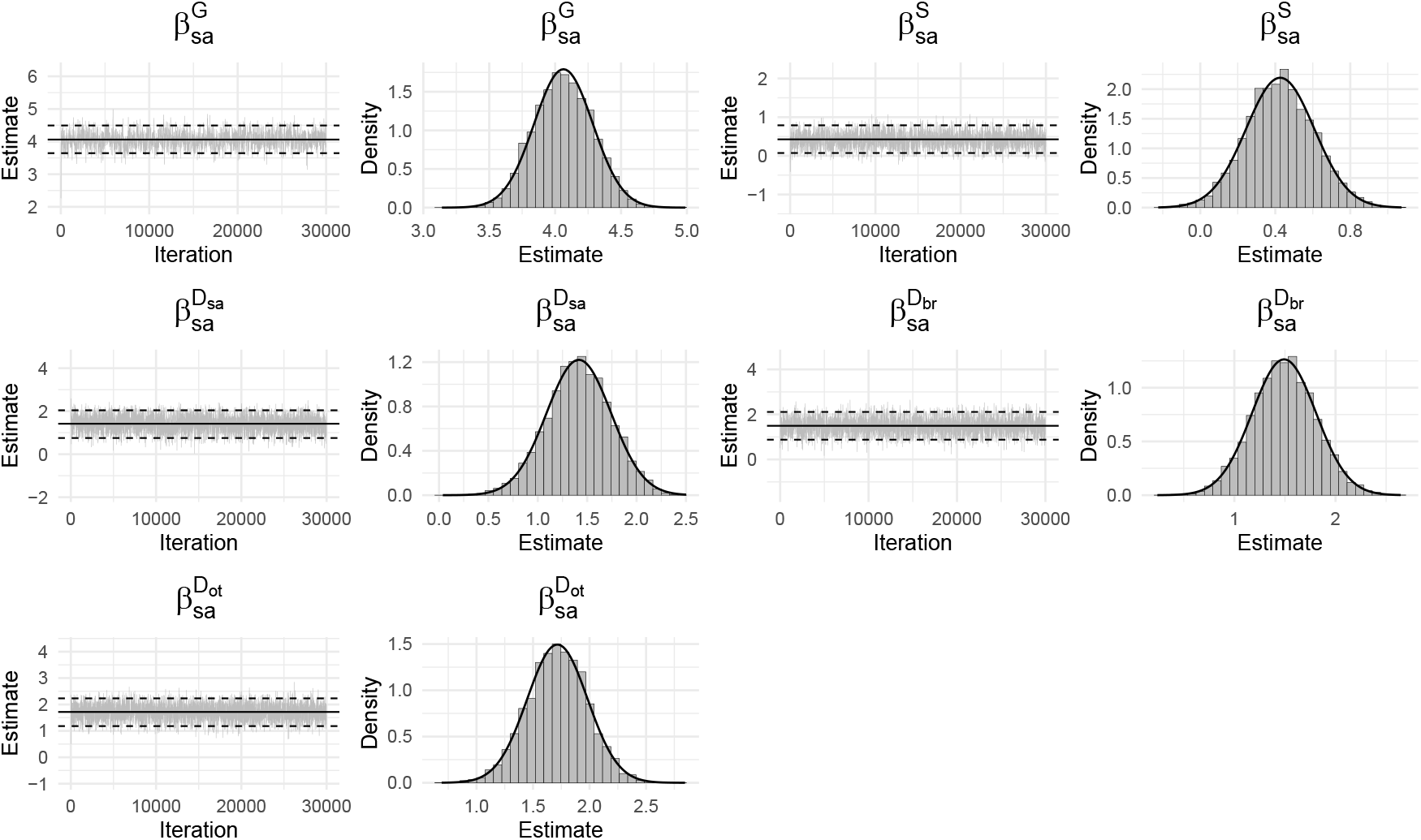
Trace plots of 30,000 posterior samples of sarcoma-specific regression coefficients ***β***_*sa*_. Solid: Estimated parameter; Dashed: 95% credible interval.

**Figure I.2:**
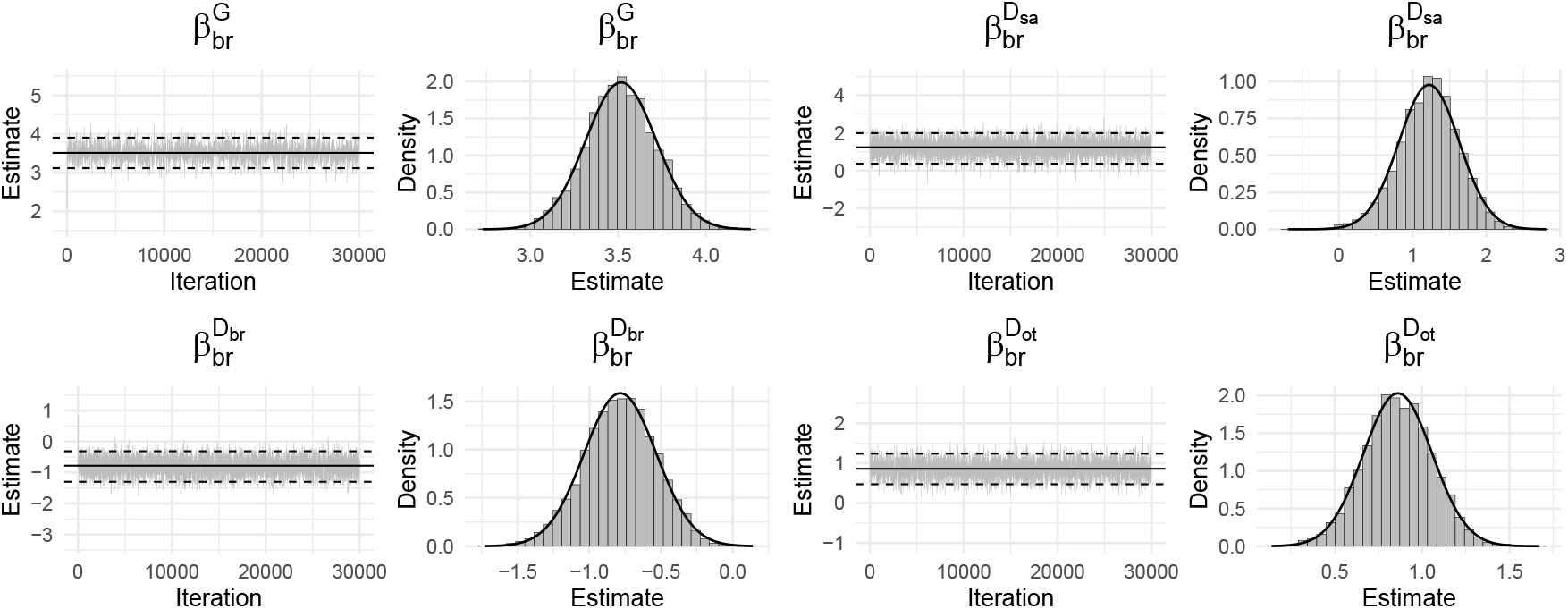
Trace plots of 30,000 posterior samples of breast-cancer-specific regression coefficients ***β***_*br*_. There is no coefficient for gender since the estimates are based on the female only. Solid: Estimated parameter; Dashed: 95% credible interval.

**Figure I.3:**
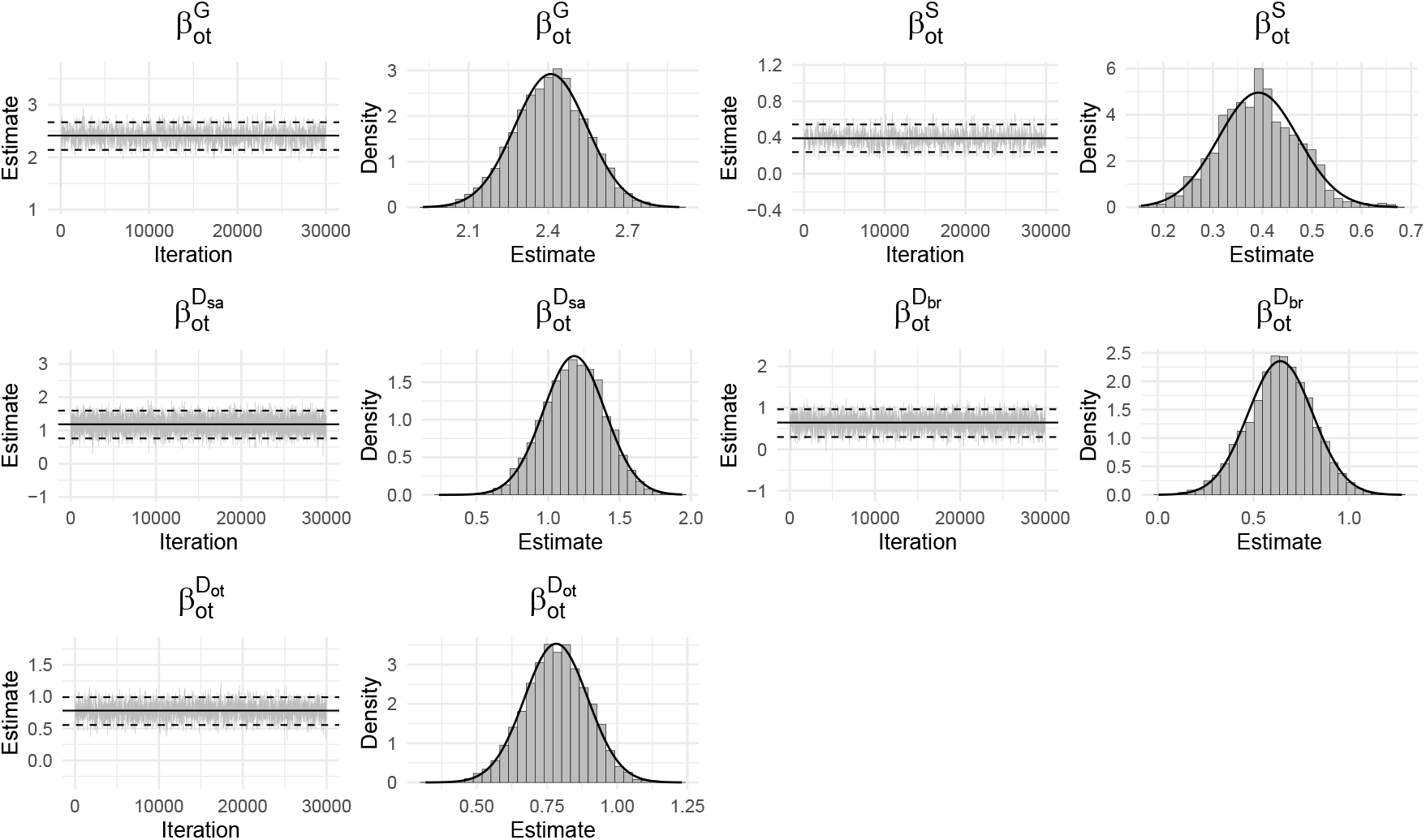
Trace plots of 30,000 posterior samples of other-cancer-specific regression coefficients ***β***_*ot*_. Solid: Estimated parameter; Dashed: 95% credible interval.

**Figure I.4:**
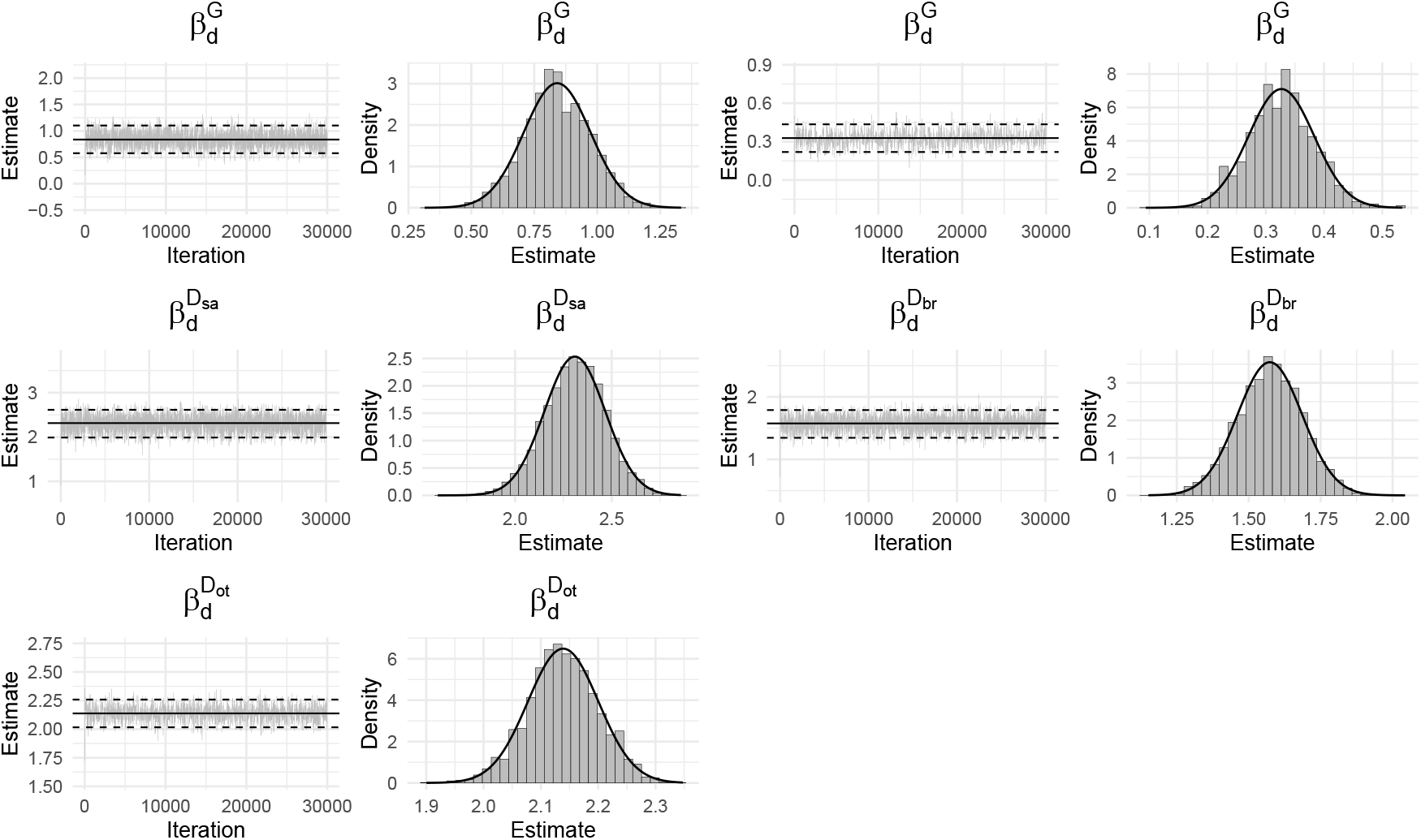
Trace plots of 30,000 posterior samples of mortality-specific regression coefficients ***β**_d_*. Solid: Estimated parameter; Dashed: 95% credible interval.

**Table I.5:** Estimated *γ* based on the last 25,000 posterior samples.

| Event | Parameter | Mean | Median | SD | 2.5% | 97.5% |
| --- | --- | --- | --- | --- | --- | --- |
| Sarcoma | $\gamma_{1,sa}$ | 0.0007 | 0.0007 | 0.0003 | 0.0002 | 0.0013 |
| | $\gamma_{2,sa}$ | 0.0010 | 0.0009 | 0.0006 | 0.0000 | 0.0024 |
| | $\gamma_{3,sa}$ | 0.0012 | 0.0010 | 0.0011 | 0.0000 | 0.0037 |
| | $\gamma_{4,sa}$ | 0.0060 | 0.0061 | 0.0026 | 0.0007 | 0.0111 |
| | $\gamma_{5,sa}$ | 0.0058 | 0.0048 | 0.0051 | 0.0000 | 0.0180 |
| Breast | $\gamma_{1,br}$ | 0.0001 | 0.0001 | 0.0002 | 0.0000 | 0.0006 |
| | $\gamma_{2,br}$ | 0.0004 | 0.0003 | 0.0005 | 0.0000 | 0.0019 |
| | $\gamma_{3,br}$ | 0.0039 | 0.0025 | 0.0044 | 0.0000 | 0.0162 |
| | $\gamma_{4,br}$ | 0.1947 | 0.1945 | 0.0222 | 0.1521 | 0.2370 |
| | $\gamma_{5,br}$ | 0.0360 | 0.0293 | 0.0320 | 0.0001 | 0.1150 |
| Other | $\gamma_{1,ot}$ | 0.0109 | 0.0108 | 0.0017 | 0.0076 | 0.0142 |
| | $\gamma_{2,ot}$ | 0.0018 | 0.0012 | 0.0019 | 0.0000 | 0.0066 |
| | $\gamma_{3,ot}$ | 0.0063 | 0.0044 | 0.0064 | 0.0000 | 0.0234 |
| | $\gamma_{4,ot}$ | 0.1391 | 0.1399 | 0.0298 | 0.0775 | 0.1965 |
| | $\gamma_{5,ot}$ | 0.4578 | 0.4540 | 0.0716 | 0.3267 | 0.6102 |
| Death | $\gamma_{1,de}$ | 0.0004 | 0.0002 | 0.0005 | 0.0000 | 0.0018 |
| | $\gamma_{2,de}$ | 0.0209 | 0.0212 | 0.0039 | 0.0127 | 0.0284 |
| | $\gamma_{3,de}$ | 0.0063 | 0.0045 | 0.0062 | 0.0000 | 0.0223 |
| | $\gamma_{4,de}$ | 0.0055 | 0.0033 | 0.0064 | 0.0000 | 0.0225 |
| | $\gamma_{5,de}$ | 1.4023 | 1.4011 | 0.0788 | 1.2532 | 1.5629 |

**Table I.6:** Estimated *ϕ* based on the last 25,000 posterior samples.

| Event | Parameter | Mean | Median | SD | 2.5% | 97.5% |
| --- | --- | --- | --- | --- | --- | --- |
| Sarcoma | $\phi_{sa}$ | 4.3845 | 4.1926 | 1.3750 | 2.2903 | 7.4901 |
| Breast | $\phi_{br}$ | 4.3634 | 4.2271 | 1.1644 | 2.4883 | 7.1033 |
| Other | $\phi_{ot}$ | 5.0060 | 4.9171 | 0.9868 | 3.3422 | 7.2996 |
| Death | $\phi_{de}$ | 5.6939 | 5.5757 | 0.9046 | 4.2380 | 7.7092 |

**Table I.7:** Estimated ***β***_*k*_, *k* ∈ {*sa,br,ot,de*}, based on the last 25,000 posterior samples. The weights *a_sa_* for sarcoma, *a_br_* for breast cancer, and *a_ot_* for other cancers are set to 4, 2, and 1 respectively.

| Event | Parameter | Mean | Median | SD | 2.5% | 97.5% |
| --- | --- | --- | --- | --- | --- | --- |
| Sarcoma | $\beta_{sa}^G$ | 3.626 | 3.624 | 0.216 | 3.199 | 4.061 |
| | $\beta_{sa}^S$ | 0.324 | 0.326 | 0.173 | -0.008 | 0.664 |
| | $\beta_{sa}^{D_{sa}}$ | 1.461 | 1.470 | 0.319 | 0.817 | 2.071 |
| | $\beta_{sa}^{D_{br}}$ | 1.444 | 1.449 | 0.308 | 0.829 | 2.050 |
| | $\beta_{sa}^{D_{ot}}$ | 1.641 | 1.647 | 0.262 | 1.118 | 2.143 |
| Breast | $\beta_{br}^G$ | 3.563 | 3.561 | 0.183 | 3.201 | 3.916 |
| | $\beta_{br}^{D_{sa}}$ | 1.246 | 1.255 | 0.380 | 0.471 | 1.982 |
| | $\beta_{br}^{D_{br}}$ | -0.794 | -0.790 | 0.248 | -1.298 | -0.323 |
| | $\beta_{br}^{D_{ot}}$ | 0.873 | 0.872 | 0.191 | 0.494 | 1.245 |
| Other | $\beta_{ot}^G$ | 2.447 | 2.448 | 0.135 | 2.183 | 2.699 |
| | $\beta_{ot}^S$ | 0.391 | 0.392 | 0.079 | 0.234 | 0.556 |
| | $\beta_{ot}^{D_{sa}}$ | 1.175 | 1.178 | 0.218 | 0.745 | 1.593 |
| | $\beta_{ot}^{D_{br}}$ | 0.633 | 0.633 | 0.168 | 0.299 | 0.961 |
| | $\beta_{ot}^{D_{ot}}$ | 0.783 | 0.782 | 0.113 | 0.563 | 1.000 |
| Death | $\beta_{de}^G$ | 0.839 | 0.843 | 0.140 | 0.555 | 1.106 |
| | $\beta_{de}^S$ | 0.328 | 0.327 | 0.057 | 0.218 | 0.441 |
| | $\beta_{de}^{D_{sa}}$ | 2.313 | 2.314 | 0.166 | 1.987 | 2.627 |
| | $\beta_{de}^{D_{br}}$ | 1.578 | 1.580 | 0.114 | 1.353 | 1.797 |
| | $\beta_{de}^{D_{ot}}$ | 2.137 | 2.134 | 0.064 | 2.012 | 2.266 |

### J. Age-at-onset Penetrance from the Family-wise Likelihood Model

**Figure J.1.**
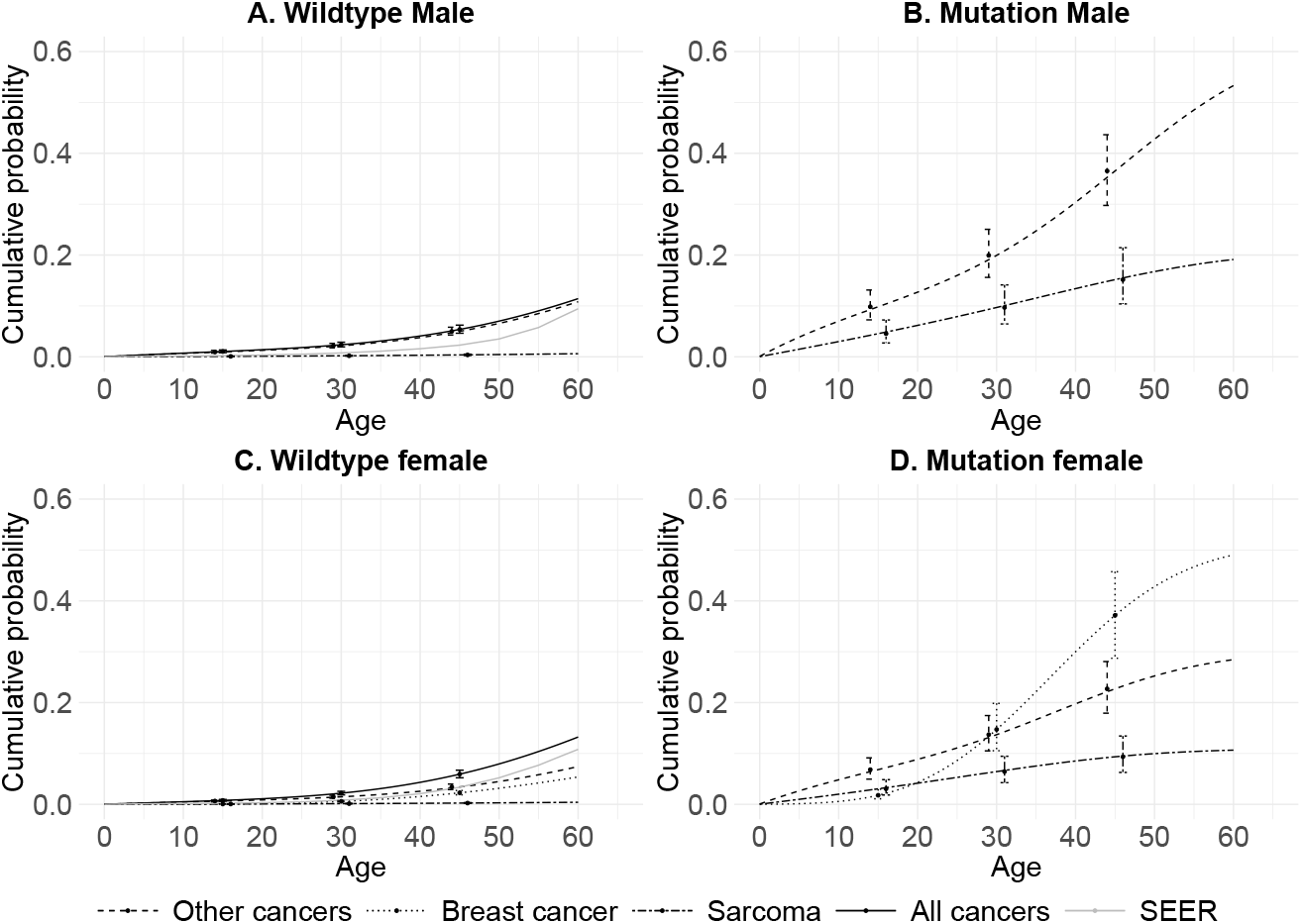
Cancer-specific age-at-onset penetrance to the first primary cancer for male and female with and without *TP53* mutation, and the corresponding 95% credible intervals at ages 15, 30, and 45. Cumulative incidence rates across all cancer types from SEER are shown for comparison.

**Figure J.2.**
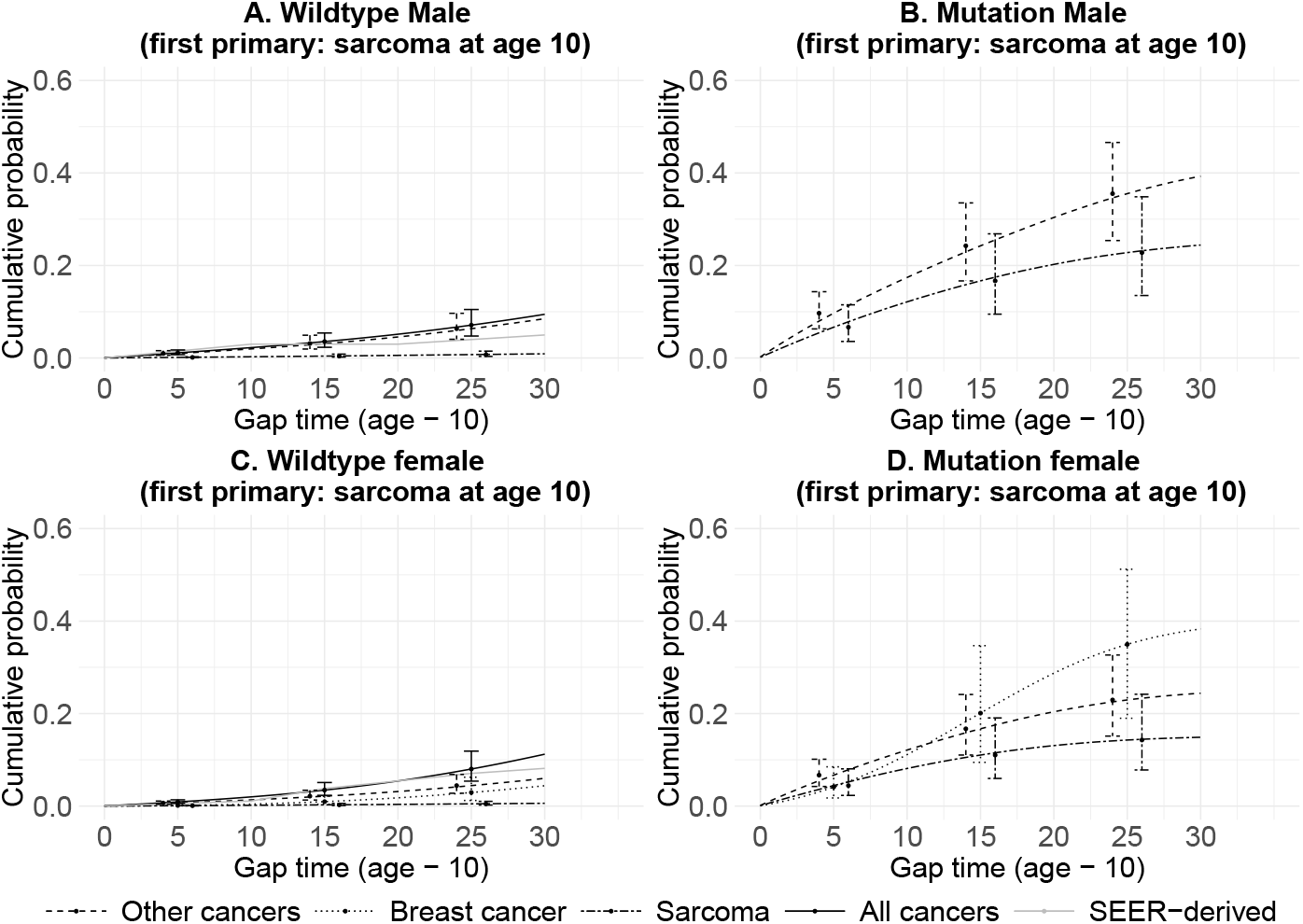
Estimates of cancer-specific age-at-onset penetrance to the second primary cancer, given the first primary of sarcoma at age 10, and the corresponding 95% credible intervals at gap times 5, 15, and 25. Cumulative incidence rates across all cancer types from SEER are shown for comparison.

**Figure J.3.**
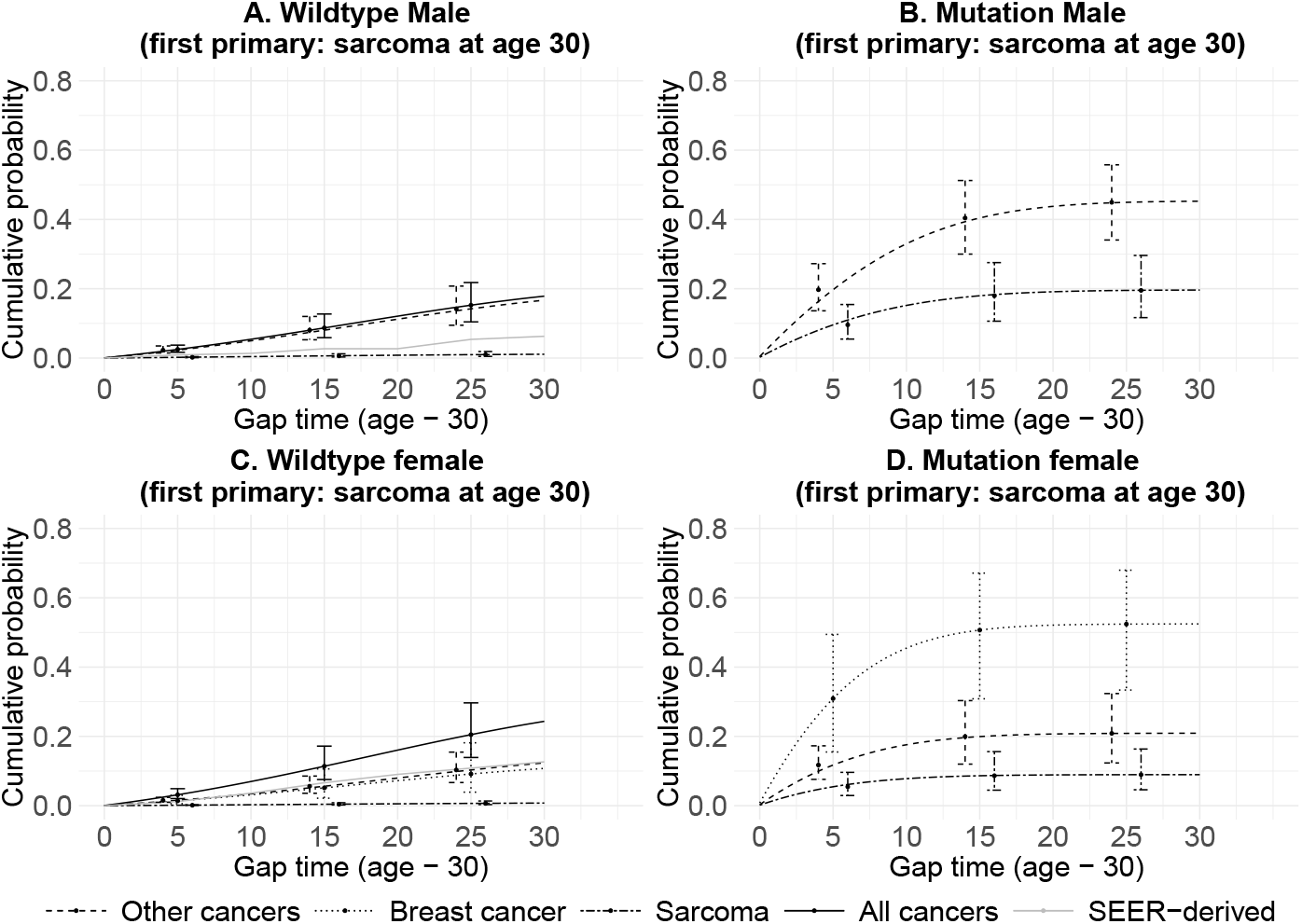
Estimates of cancer-specific age-at-onset penetrance to the second primary cancer, given the first primary of sarcoma at age 30, and the corresponding 95% credible intervals at gap times 5, 15, and 25. Cumulative incidence rates across all cancer types from SEER are shown for comparison.

**Figure J.4.**
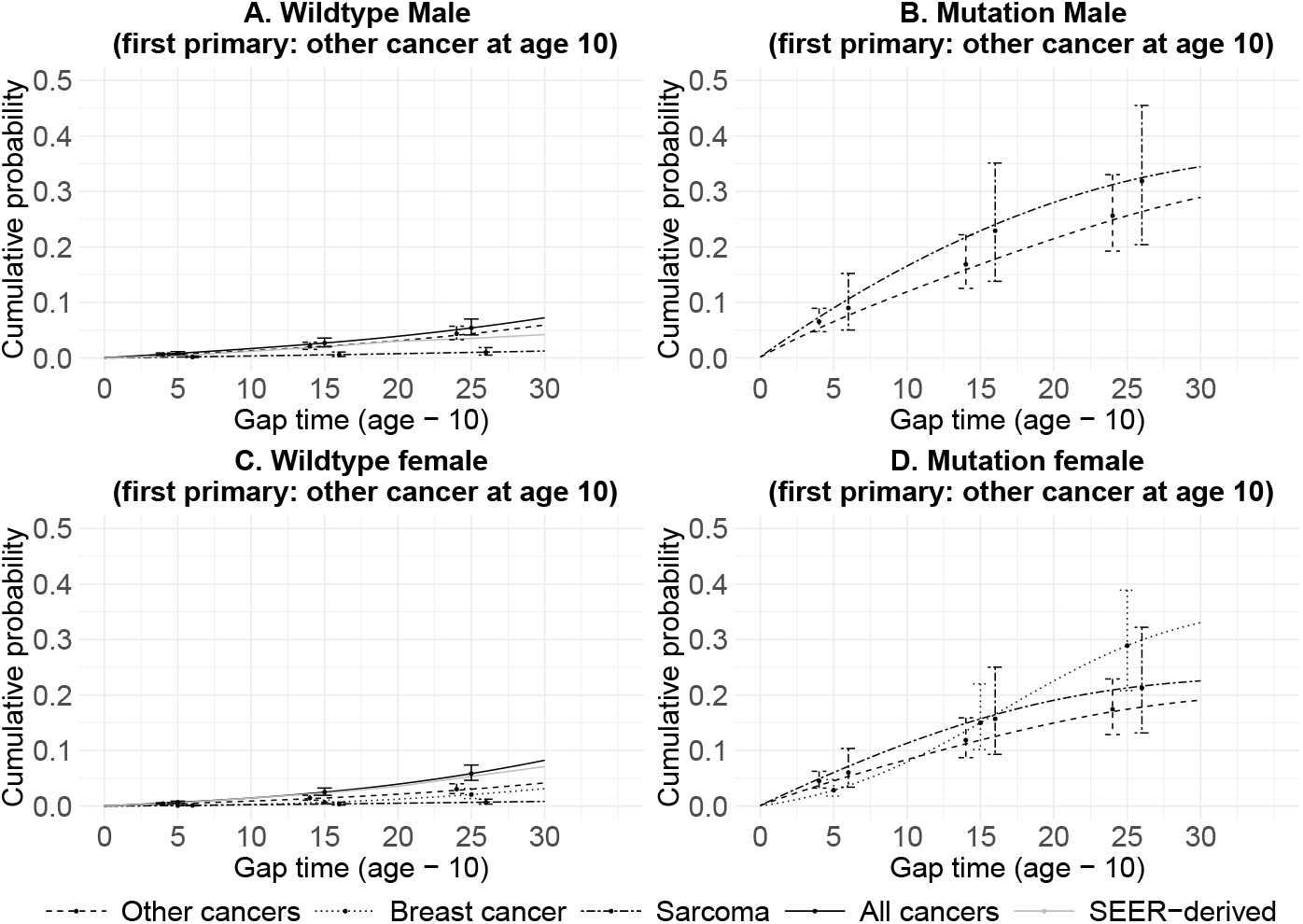
Estimates of cancer-specific age-at-onset penetrance to the second primary cancer, given the first primary other than breast cancer and sarcoma at age 10, and the corresponding 95% credible intervals at gap times 5, 15, and 25. Cumulative incidence rates across all cancer types from SEER are shown for comparison.

**Figure J.5.**
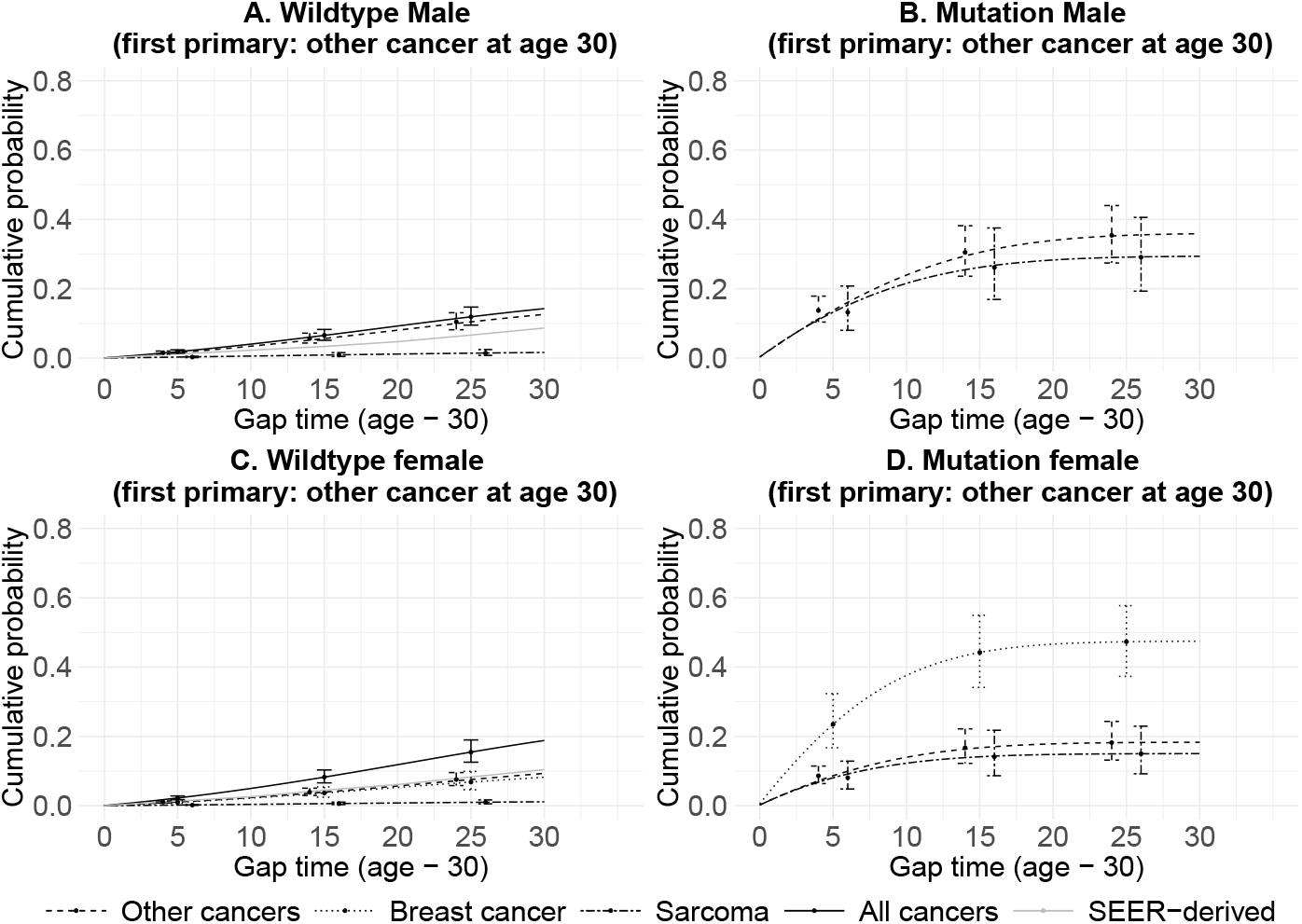
Estimates of cancer-specific age-at-onset penetrance to the second primary cancer, given the first primary other than breast cancer and sarcoma at age 30, and the corresponding 95% credible intervals at gap times 5, 15, and 25. Cumulative incidence rates across all cancer types from SEER are shown for comparison.

### K. Analyses Results from the Validation Study of the Family-wise Likelihood Model

**Figure K.1:**
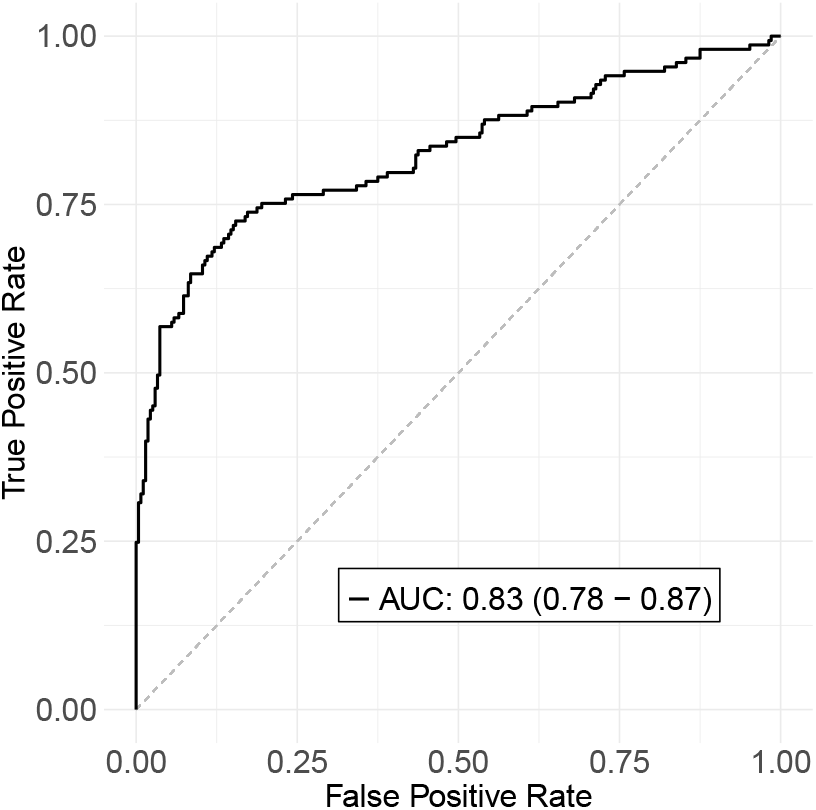
ROC curve, along with the AUC and 95% bootstrapped confidence interval, for prediction of *TP53* mutation in the validation dataset. Sample size: n(wildtype) = 272, n(mutation) = 153.

**Figure K.2:**
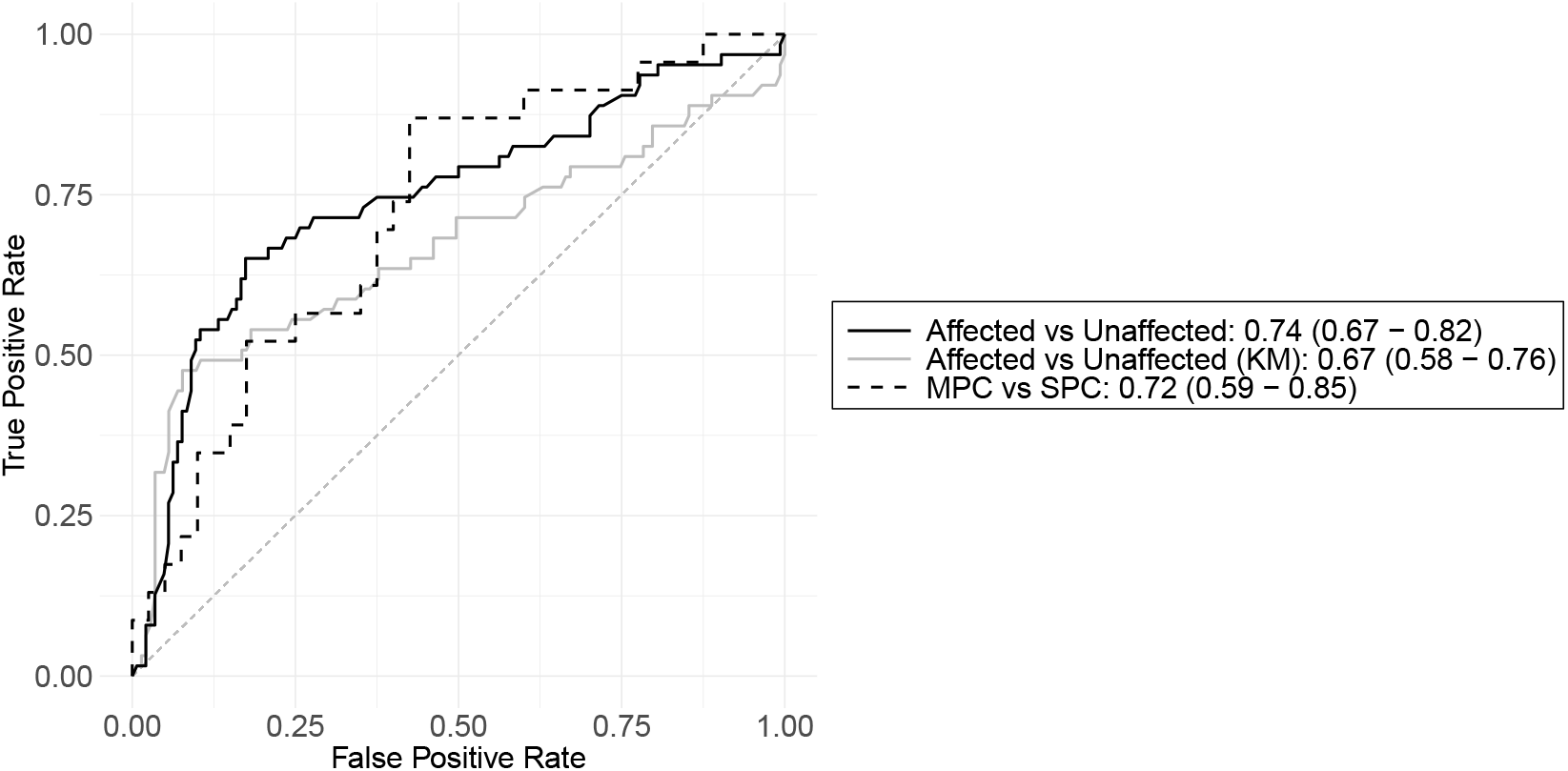
ROC curves, along with the AUCs and their 95% bootstrapped confidence intervals, for prediction of the number of primary cancer in the validation dataset. FPC = first primary cancer, SPC = second primary cancer. Sample size: n(unaffected) = 144, n(FPC) = 38, n(SPC) = 23.

**Figure K.3:**
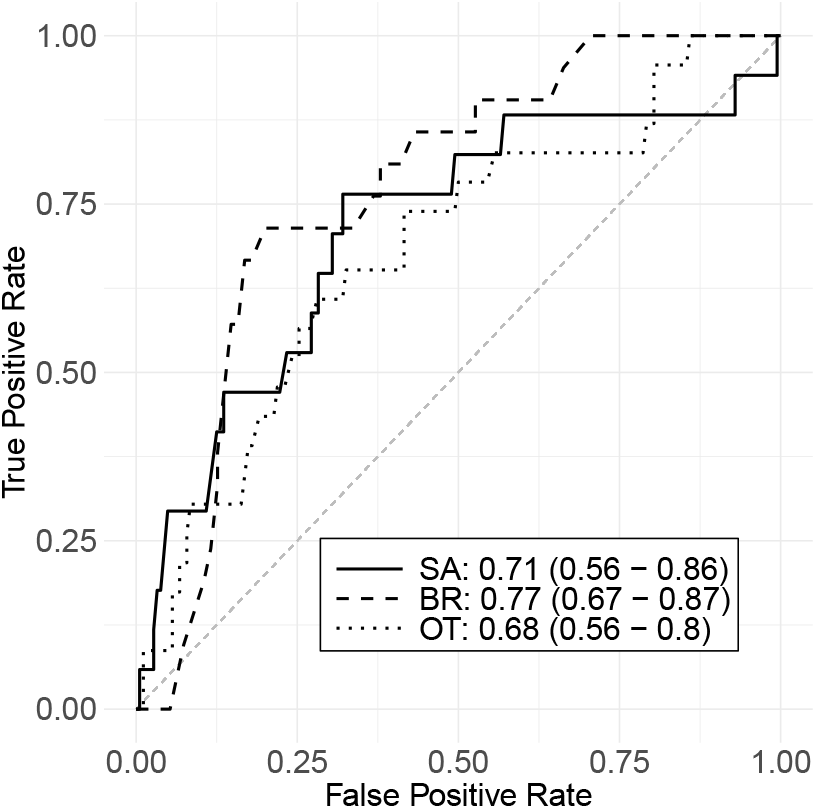
ROC curves, along with the AUCs and their 95% bootstrapped confidence intervals, for cancer-specific prediction of FPC in the validation dataset. SA = sarcoma, BR = breast cancer, OT = other cancers combined. Sample size: n(SA) = 17, n(BR) = 21, n(OT) = 23.

